# Myeloid-mesenchymal crosstalk drives Arg1-dependent profibrotic metabolism via ornithine in lung fibrosis

**DOI:** 10.1101/2023.09.06.556606

**Authors:** Preeti Yadav, Javier Gómez Ortega, Prerna Dabral, Whitney Tamaki, Charles Chien, Kai-chun Chang, Nivedita Biswas, Sixuan Pan, Julia Nilsson, Xiaoyang Yin, Aritra Bhattacharyya, Kaveh Boostanpour, Tanay Jujaray, Jasper Wang, Tatsuya Tsukui, Chris Molina, Vincent Auyeung, Dean Sheppard, Baosheng Li, Mazharul Maishan, Hiroki Taenaka, Michael A. Matthay, Rieko Muramatsu, Lenka Maliskova, Arnab Ghosh, Walter L. Eckalbar, Ari B. Molofsky, Stanley J. Tamaki, Trever Bivona, Adam R. Abate, Allon Wagner, Satish K. Pillai, Paul J. Wolters, Kevin M. Tharp, Mallar Bhattacharya

## Abstract

Idiopathic pulmonary fibrosis (IPF) is a disease of progressive lung remodeling and collagen deposition that leads to respiratory failure. Myeloid cells are abundant in IPF lung and in murine lung fibrosis, but their functional effects are incompletely understood. Using mouse and human lung models, we show that ornithine produced by myeloid cells expressing Arg1 serves as a substrate for proline and collagen synthesis by lung fibroblasts. The predominant Arg1-expressing myeloid cells in mouse lung were macrophages, but in IPF lung, high-dimensional imaging revealed ARG1 to be expressed mainly in neutrophils. Arg1 inhibition suppressed both ornithine levels and collagen expression in cultured, precision-cut IPF lung slices and in murine lung fibrosis. These results were confirmed in macrophage-specific Arg1 KO mice. Furthermore, we find that this pathway is regulated by cell-to-cell crosstalk, starting with purinergic signaling: Fibroblast eATP receptor P2rx4 was necessary for fibroblast IL-6 expression, which in turn was necessary for Arg1 expression by myeloid cells. Taken together, our findings define an immune-mesenchymal circuit that governs profibrotic metabolism in lung fibrosis.

## Introduction

Pathologic fibrosis is a destructive response to tissue injury that results from the deposition of excess fibrillar collagen by activated fibroblasts. In idiopathic pulmonary fibrosis (IPF), it is believed that dysfunction of the epithelium leads to the recruitment and activation of multiple cell types, including most importantly fibroblasts and myeloid cells (1). These cells have been found to be in close proximity in the fibrotic niche (2, 3), but it remains unclear how they interact to induce the fibrotic phenotype. One approach is to define and test the functional role of diffusible factors that have been detected in fibrosis and that might subserve crosstalk between the various cell types leading to collagen production by fibroblasts.

We recently considered the role of extracellular ATP (eATP), a damage-associated molecular pattern (DAMP) molecule that is extensively present in the IPF lung, as revealed in the bronchoalveolar lavage from patients for example (4). eATP’s role in fibrosis remains incompletely understood.

Although eATP activates receptor-mediated transcriptional responses by ligating multiple purinergic receptors, in lung fibroblasts we reported P2rx4 as the critical purinergic receptor: Fibroblast-specific deletion of the eATP receptor P2rx4 decreased lung fibrosis in mice (5). However, the paracrine signaling effects of this pathway are not known.

Here we explore the effect of P2rx4 signaling on cell-to-cell crosstalk in fibrosis. Recent studies have revealed a heterogeneity of fibroblast states following injury, and inflammatory fibroblasts have been detected as a distinct subset of mesenchymal cells that express secreted inflammatory factors such as chemokines and cytokines (6–8). How these signals from inflammatory fibroblasts promote fibrosis is unknown. We hypothesized that eATP-P2rx4 signaling may stimulate expression of these diffusible signals from fibroblasts after injury, initiating profibrotic crosstalk with other cells in the fibrotic niche.

Studies with murine macrophage-fibroblast cocultures revealed that macrophage Arg1 expression was dependent on P2rx4 signaling in fibroblasts. Arg1+ macrophages were increased after bleomycin-induced lung fibrosis in mice; in IPF lung, ARG1+ cells were also increased, in the neutrophil compartment. In both murine and human lung systems, Arg1 inhibition decreased lung fibrosis.

Furthermore, Arg1 was induced by fibroblast expression of the cytokine IL-6 in a P2rx4-dependent manner. Mechanistically, Arg1 drove fibroblast collagen synthesis via the production of the noncoding amino acid ornithine, which served as a precursor for the collagen building block proline. These findings indicate a critical profibrotic role for a metabolic pathway arising from cellular crosstalk in the development of lung fibrosis.

## Results

### Arg1+ cells localize to the fibrotic niche

We recently reported that mice with deletion of the eATP receptor P2rx4 in fibroblasts were protected from bleomycin injury-induced lung fibrosis (5). To screen for macrophage genes regulated by fibroblast eATP-P2rx4 signaling, we used an *in vitro* system: Either WT or P2rx4 KO fibroblasts were cocultured with WT lung macrophages (**Supplemental Figure 1A**), followed by single cell RNAseq. Both cell types were isolated from murine lungs 7 days after injury to approximate cellular states during the early fibrotic period, defined by increased fibroblast expression of Col1a1 and Col3a1 (**Supplemental Figure 1B**). Fibroblasts were used directly, whereas macrophages were treated for 48 hours in culture with Csf1 to maintain macrophage identity prior to coculture. After 5 days of coculture, cells were submitted for scRNAseq.

The most downregulated macrophage gene when fibroblast P2rx4 was deleted was Arg1, a marker of alternatively activated macrophages (**Figure 1A**). Analysis of lung macrophages in a published scRNAseq time course following bleomycin lung injury (9) showed maximal *Arg1* in the “C2”, transitional macrophage compartment that we previously demonstrated was localized to the fibrotic niche after injury(2) (**Figure 1B**). C2 cells expressed higher levels of Arg1 and also the “Fab5” marker genes that were recently found to be associated with a core pro-fibrotic macrophage program (10) (**Supplemental Figure 1C-E**), and they were transcriptomically similar to recruited and interstitial macrophages (11) (**Supplemental Figure 1F**). Interestingly, the macrophages from our cocultures also expressed higher levels of interstitial macrophage than alveolar macrophage genes (**Supplemental Figure 1G**). To determine whether Arg1+ cells localized to areas of fibrosis, we injured mice expressing reporter alleles for Arg1 and for the fibroblast marker Col1a1. Confocal imaging revealed that Arg1+ cells were in proximity to clusters of activated fibroblasts in the fibrotic niche (**Figure 1C**).

**Figure 1:**
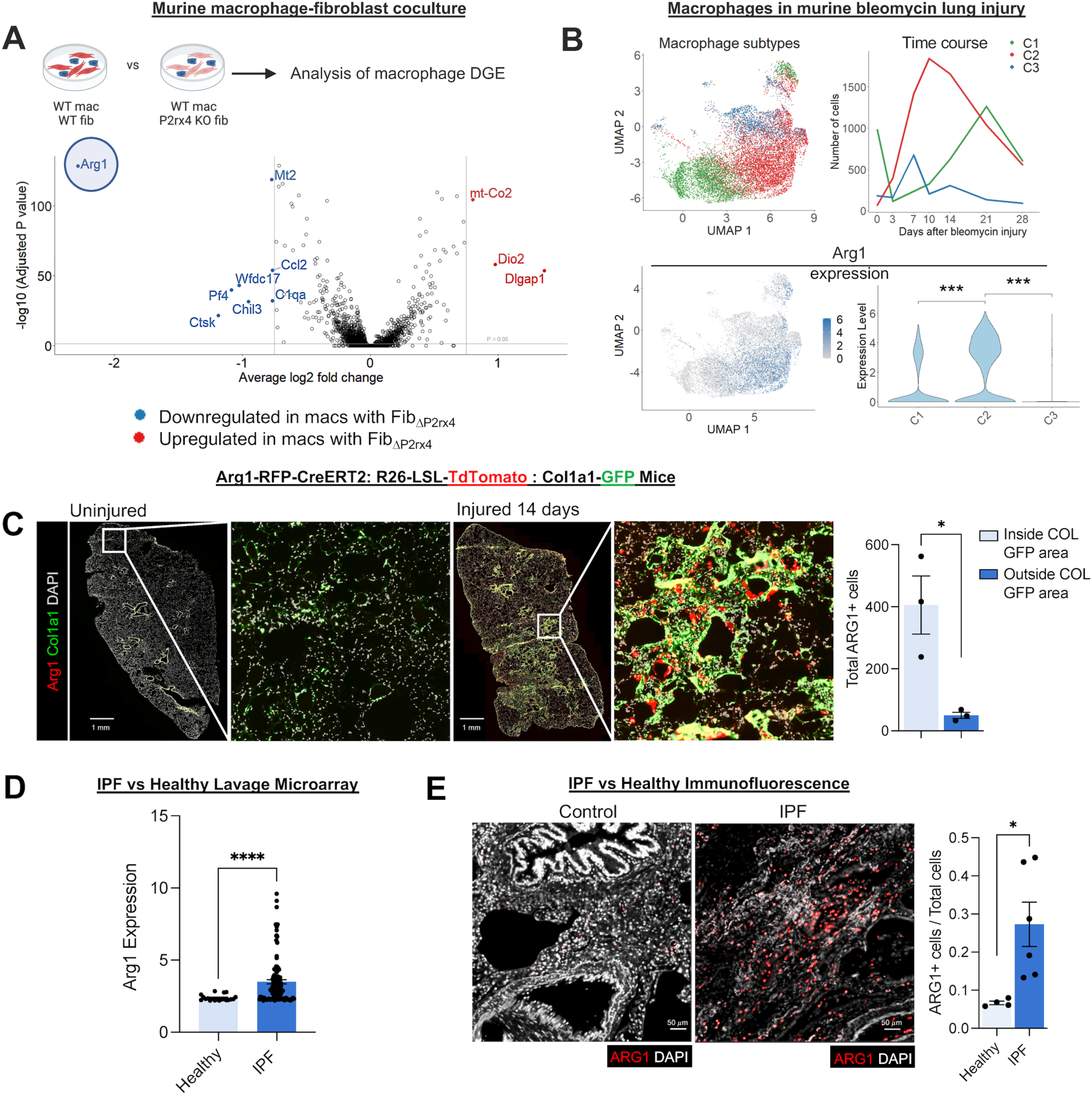
**Arginase 1-expressing cells localize to the fibrotic niche in murine and human lung.** A) Volcano plot for macrophages from scRNAseq of macrophage-fibroblast cocultures for WT macrophages and fibroblasts versus WT macrophages cocultured with P2rx4 KO (*Pdgfrb-Cre: P2rx4 f/f*) fibroblasts. Significance was determined by Wilcoxon Rank Sum test corrected for multiple comparisons by Bonferroni method. Color-labeled genes had p<0.05 and absolute value of log2 fold change >0.75. Data represent n=2 separate cocultures. B) Analysis of macrophages from bleomycin lung injury (data reanalyzed from^7^). Top left: Annotation of macrophages from multiple time points, according to C1, C2, or C3 macrophage annotation (as described in (7): C1=alveolar macrophages; C2=transitional monocyte-derived macrophages; C3=monocyte derived macrophages). Top right: Proportions of C1, C2, and C3 across time. Bottom left: Feature plot of Arg1 expression. Bottom right: Violin plot of Arg1 expression according to cluster (C1, C2, and C3). ***p<0.001 by Student’s t-test. C) Immunofluorescence imaging of tamoxifen-induced Arg1-RFP-CreERT2: LSL-tdTomato-Col1a1-GFP mice at 14 days after bleomycin injury. Quantification shows total Arg1+ cells inside and outside contiguous Col1a1-GFP+ areas. *p<0.05 by Student’s t-test. +/- SEM. D) *ARG1* expression by microarray analysis of gene expression of BAL cells from healthy (20 patients) and IPF (112 patients) from GSE70867(12). ****p<0.0001 by Mann-Whitney test. +/- SEM. E) Representative immunofluorescence of human healthy control and IPF lung sections (n=4 and n=6, respectively) for ARG1. The quantification shows the proportion of ARG1+ cells (i.e., ARG1+ / total DAPI count) for each sample. *p<0.05 by Student’s t-test. +/- SEM.

To further profile Arg1 expression in myeloid cell subsets in mice, we performed flow cytometry with Arg1 reporter mice. This analysis confirmed that Arg1 was expressed 10 days after bleomycin injury in macrophages expressing CD11b and CD64, markers consistent with monocyte-derived macrophages (**Supplemental Figure 2A-C**). Canonical neutrophils expressing Ly6G did not express Arg1, which we confirmed by immunofluorescence for the neutrophil marker S100A8 (**Supplemental Figure 2D**). However, we did note emergence of a small myeloid population of Arg1+ cells expressing Ly6G that also expressed CD64 and CD11c, though this population was about tenfold smaller in number than moMacs (**Supplemental Figure 2E**). Taken together, these data confirm expression of Arg1 in myeloid cells in the mouse lung fibrotic niche.

To determine the relevance of ARG1 to IPF, we first interrogated a published dataset of BAL cell gene expression by microarray (12). This analysis confirmed higher ARG1 expression in IPF compared to healthy controls (**Figure 1D**). We then found increased ARG1+ cells by immunofluorescence of IPF lung explants acquired at the time of clinical transplantation compared with deceased donor lungs not known to have lung disease (**Figure 1E; Supplemental Table 1**). In human lung, ARG1 has been found to be expressed in neutrophils, based on data from lung cancer specimens (13–15). However, deep phenotyping in IPF lung tissue of ARG1-expressing cell types has not been undertaken. Therefore, to comprehensively profile the cell types expressing ARG1 in IPF, we performed multiplexed ion beam imaging (MIBI) (16–19), an antibody-based imaging technique in which isotopically pure elemental metal reporters are conjugated to antibodies and detected spatially in combination with mass spectrometry. We applied MIBI to sections from 5 IPF explanted lungs using anti-ARG1 antibody and 33 other antibodies, selecting markers primarily for the purpose of defining major immune cell types. These data showed that ARG1+ cells predominantly expressed the neutrophil-specific markers CD66B and MPO but not the macrophage markers CD68, CD206, and CD163 (**Figure 2A-C; Supplemental Table 2**). Importantly, neutrophils have been associated with increased mortality in IPF (20–23). However, their function in fibrogenesis has not been well studied.

**Figure 2:**
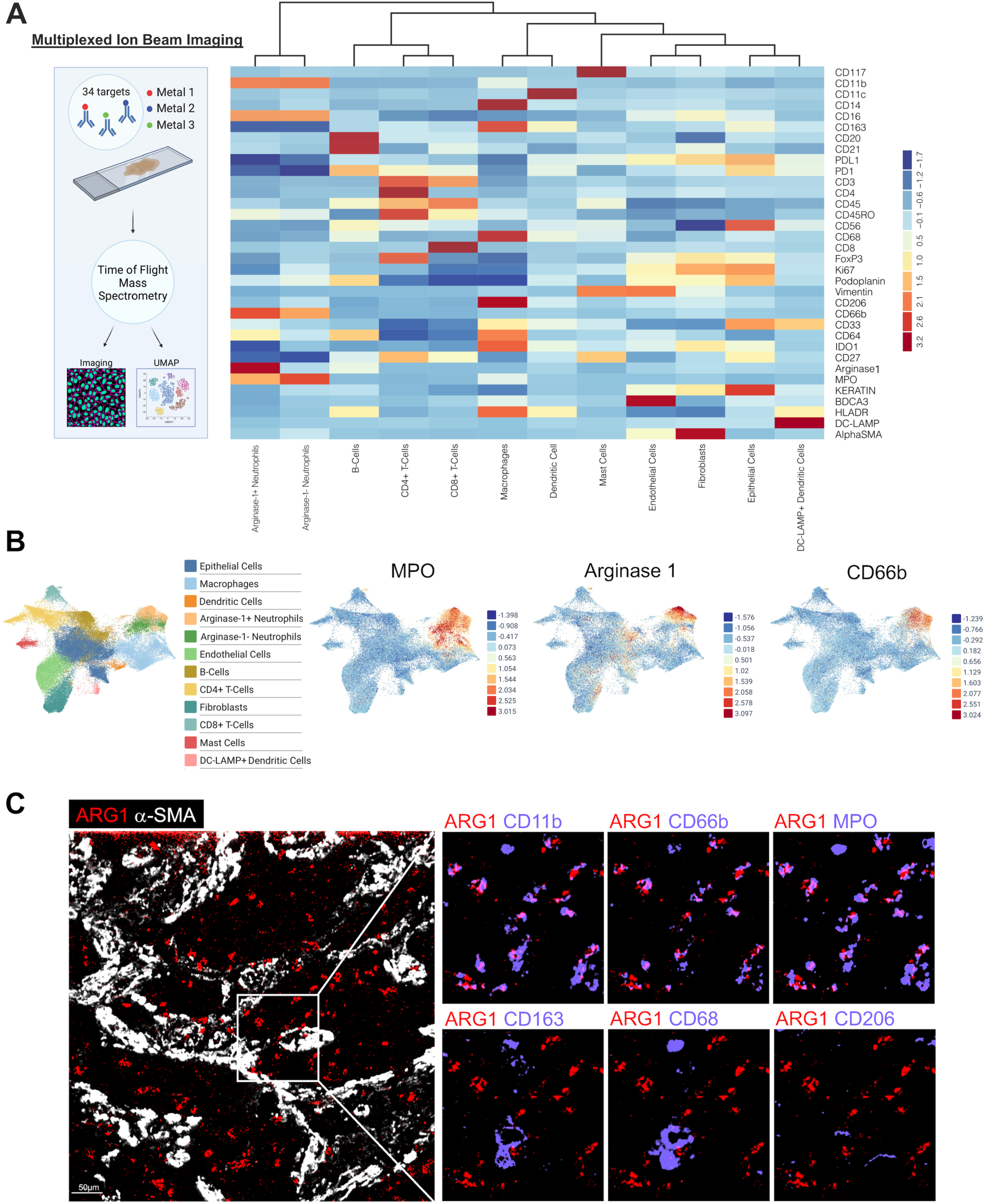
**Arginase 1 is expressed predominantly by neutrophils in IPF.** A) Schematic of multiplexed ion beam imaging (MIBI) and heatmap of scaled marker expression from MIBI of sections from n=5 IPF explanted lungs. B) MIBI UMAPs by cluster and individual markers. C) MIBI images for individual markers within a representative field of view of IPF lung.

### Arg1 regulates availability of ornithine, a pro-fibrotic substrate

To understand Arg1’s function, we considered first that ornithine generated from arginine by Arg1 can serve as a substrate for the synthesis of proline (24), a major constituent of collagen. Remarkably, we found that ornithine was markedly increased both in mouse lung 14 days after bleomycin injury compared to steady state and in IPF lung lysates compared to healthy controls (**Figure 3A**). To test the effects of Arg1 and ornithine in fibrosis, we first used the small-molecule Arg1 inhibitor CB-1158 (25). We confirmed the on-target effect of CB-1158 by measuring lung ornithine after treatment: In both bleomycin-injured mice and precision-cut lung slices (PCLS) from explanted IPF lungs, the inhibitor decreased ornithine levels (**Figure 3B**). Next, we treated bleomycin-injured mice with CB-1158 during the fibrotic period. Importantly, we found that lung fibrosis was significantly decreased and body weight recovery improved with Arg1 inhibition (**Figure 3C; Supplemental Figure 2F**). We then prepared PCLS from explanted IPF lungs and treated them with CB-1158 for 24 hours, followed by immunoblot for COL1A1 in the RIPA buffer-soluble fraction representing the more soluble, newly synthesized collagen (26). COL1A1 expression measured by immunoblot of lung slice lysates was decreased by CB-1158 treatment compared to untreated controls, indicating suppression of collagen synthesis with ARG1 inhibition (**Figure 3D**). Retention of cells expressing ARG1 and neutrophil markers MPO and CD66B was confirmed in PCLS by immunofluorescence (**Supplemental Figure 3A**). Finally, taking a genetic approach with macrophage-specific Arg1 KO mice in which Cre-mediated deletion efficiency was approximately 55% (**Supplementary** Figure 3B), we found that lung fibrosis was significantly decreased after injury in KO mice compared to controls (**Figure 3E**). Taken together, these data indicate that Arg1 determines the pathologic accumulation of collagen in lung fibrosis.

**Figure 3:**
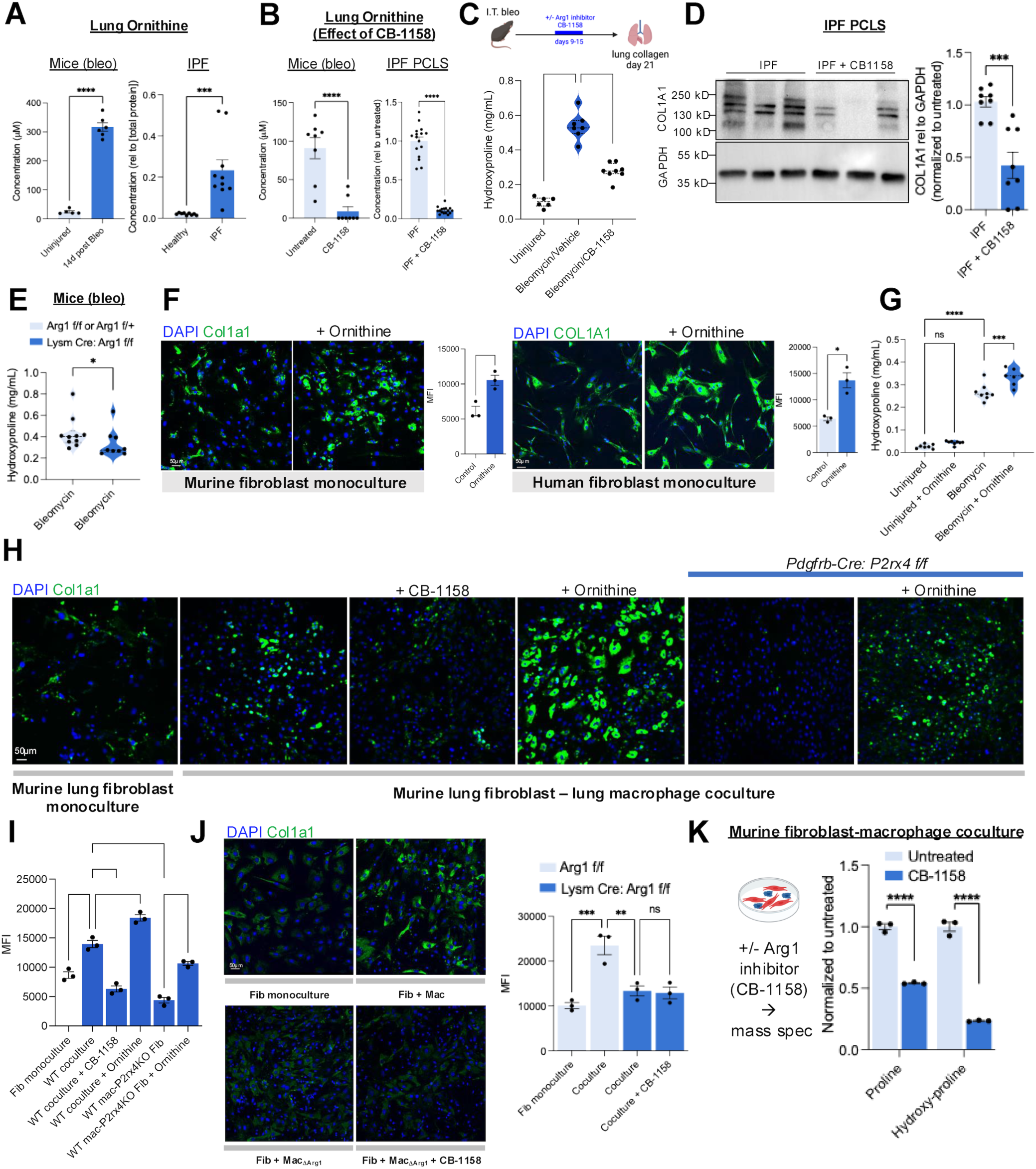
**Arginase 1 regulates lung collagen via ornithine production.** A) Ornithine concentration measured in lysates of mouse lung (N=5 uninjured and N=6 injured mice), healthy human donor lung (N=9) and patient IPF lung explants (N=10). ****p<0.0001, ***p<0.001 by Student’s t-test. +/- SEM. B) Ornithine concentration with and without CB-1158 treatment of lungs of bleomycin-injured mice (N=8 in each condition) and cultured IPF PCLS (N=16 slices from a total of 3 patients). ****p<0.0001 by Student’s t-test. +/- SEM. C) Hydroxyproline assay for collagen content of lungs from injured and uninjured WT mice treated with or without Arg1 inhibitor CB-1158. Mice were dosed with 100 mg/kg of CB-1158 twice daily during days 9-15 post bleomycin. N=6, 7, 8 mice, left to right. ****p<0.0001 by 1-way ANOVA with post hoc Sidak’s multiple comparisons tests. Median, upper, and lower quartiles indicated by dashed lines. D) Representative immunoblot for COL1A1 from lysates of precision-cut IPF lung slices cultured for 24 hours with or without Arg1 inhibitor CB-1158. Quantitation is for slices from a total of n=3 patients. ***p<0.001 by Student’s t-test. +/- SEM. E) Hydroxyproline assay for collagen content of lungs from mice 21 days after bleomycin injury. N=10 and 9 mice, left to right. *p<0.05 by Mann-Whitney test. Median, upper, and lower quartiles indicated by dashed lines. F) *Left:* Col1a1 immunofluorescence of monocultured mouse lung fibroblasts with or without ornithine treatment. Quantification is for n=3 separate cultures each. *p<0.05 by Student’s t-test. *Right:* COL1A1 immunofluorescence of monocultured human lung fibroblasts with or without ornithine treatment. Quantification is for n=3 separate cultures each. *p<0.05 by Student’s t-test. +/- SEM. G) Hydroxyproline assay for collagen content of lungs from injured WT mice treated with or without ornithine (2 g/kg) twice daily by ornithine gavage during days 7 through 20. N=8, 7 mice, left to right. ***p<0.001, ****p<0.0001 by 1-way ANOVA with post hoc Sidak’s multiple comparisons tests. Median, upper, and lower quartiles indicated by dashed lines. H) Representative samples of Col1a1 immunofluorescence of mouse lung macrophage-fibroblast cocultures from WT or fibroblast-specific P2rx4 KO (Pdgfrb-Cre: P2rx4 f/f) mice treated with or without CB-1158 or ornithine. I) Quantitation of Mean Fluorescence Intensity (MFI) from (H). N=3 biological replicates per condition. ****p<0.0001 by 1-way ANOVA with post hoc Sidak’s multiple comparisons tests. +/- SEM. J) Representative samples of Col1a1 immunofluorescence of mouse lung macrophage-fibroblast cocultures treated with or without CB-1158. “Mac_λ1Arg1_” indicates macrophages isolated from Lysm-Cre: Arg1 f/f mice, whereas “Mac” indicates Arg1 f/f controls. **p<0.01, ***p<0.0001 by 1-way ANOVA with post hoc Sidak’s multiple comparisons tests. +/- SEM. K) Relative quantities of proline and hydroxy-proline in lysates of murine primary lung fibroblasts isolated after coculture with macrophages, with or without CB-1158 inhibitor treatment, quantified by liquid chromatography-mass spectrometry. N=3 biological replicates for macrophages and fibroblasts, respectively. ****p<0.0001 by 1-way ANOVA with post hoc Sidak’s multiple comparisons tests. +/- SEM.

We then tested the effect of ornithine on collagen expression. First, we found that exogenous ornithine increased fibroblast collagen expression by immunofluorescence in monocultured fibroblasts from both mouse and human lung (**Figure 3F**). To test the profibrotic potential of ornithine *in vivo*, we treated mice with ornithine by oral gavage, either at steady state or during the fibrotic phase after bleomycin injury. Ornithine treatment increased lung collagen production in the setting of injury but not at steady state, consistent with increased cellular demand for collagen synthetic precursors including proline during fibrogenesis (**Figure 3G**). We then found that macrophage-fibroblast coculture increased collagen expression compared to fibroblast monoculture, an effect that could be blocked with CB-1158 or fibroblast P2rx4 KO and enhanced with ornithine (**Figure 3H-I**). Furthermore, ornithine could rescue the CB-1158-dependent decreased collagen expression in cocultures (**Supplemental Figure 3C**).

Finally, fibroblast coculture with Arg1-specific KO macrophages was associated with decreased collagen expression compared with WT macrophages, an effect that was not significantly augmented with CB-1158 (**Figure 3J**). Taken together, these results indicate a direct profibrotic effect of ornithine *in vitro* and *in vivo*.

Ornithine is converted by ornithine aminotransferase (OAT) to glutamate-5-semialdehyde, which spontaneously cyclizes to pyrroline-5-carboxylate and is then reduced by P5C reductases to proline.

Notably, OAT inhibition decreased coculture-dependent augmentation of collagen expression (**Supplemental Figure 3C**). Interestingly, coculture also increased α-SMA expression, but this effect was not augmented by ornithine or suppressed by OAT inhibition (**Supplemental Figure 3D)**. These data indicate a dependence of collagen synthesis on Arg1 function and ornithine metabolism. Finally, we directly tested the hypothesis that Arg1-mediated ornithine production was necessary for increasing fibroblast proline content, using the approach of macrophage-fibroblast coculture followed by isolation of fibroblasts for LC-MS to detect proline. Consistent with our hypothesis, Arg1 inhibition with CB-1158 significantly decreased fibroblast proline content in cocultures (**Figure 3K**). Taken together, these results suggest that macrophage Arg1 drives fibrosis by producing ornithine, which serves as a substrate for fibroblast proline synthesis, augmenting collagen expression.

### Arg1 is regulated by IL-6

To determine how macrophage Arg1 expression is regulated by signaling within the fibrotic niche, we considered that our previously described coculture data (**Figure 1A**) indicated that P2rx4 signaling in fibroblasts was necessary for Arg1 expression in cocultured macrophages. Therefore, we performed analysis of ligand-receptor interactions via CellChat (27) and Ingenuity Pathways Analysis. This analysis revealed suppression of IL-6 signaling in the P2rx4 KO condition (**Figure 4A**). We also found that P2rx4 KO fibroblasts expressed less IL-6 mRNA compared to WT (**Figure 4B**). IL-6 is an important profibrotic factor and is a therapeutic target in clinical lung fibrosis (28–31). We first confirmed an increase of Arg1 in cultured mouse lung macrophages treated with IL-6 (**Figure 4C**). *In vivo*, we noted that IL-6 KO mice had decreased macrophage numbers in the lung, consistent with a chemotactic effect of IL-6 (**Supplemental Figure 3E**). Nonetheless, even accounting for this lower cell count, Arg1 lavage levels were disproportionately decreased in IL-6 KO mice, as indicated by bronchoalveolar lavage Arg1 concentrations normalized to CD11b+CD64+ macrophage count and by qPCR measurement of *Arg1* relative to *Gapdh* in flow-sorted CD11b+CD64+ macrophages (**Figure 4D**). Of note, IL-6 KO mice had decreased lung fibrosis after bleomycin lung injury (**Supplemental Figure 3F**). We then analyzed scRNAseq time course data for multiple lung cell types and found that IL-6 was indeed highly expressed in fibroblasts relative to other cell types (**Figure 4E**). Furthermore, IL-6 was notably expressed in the emergent, disease-associated inflammatory and fibrotic fibroblasts observed in recent time course data (8) (**Figure 4F**).

**Figure 4:**
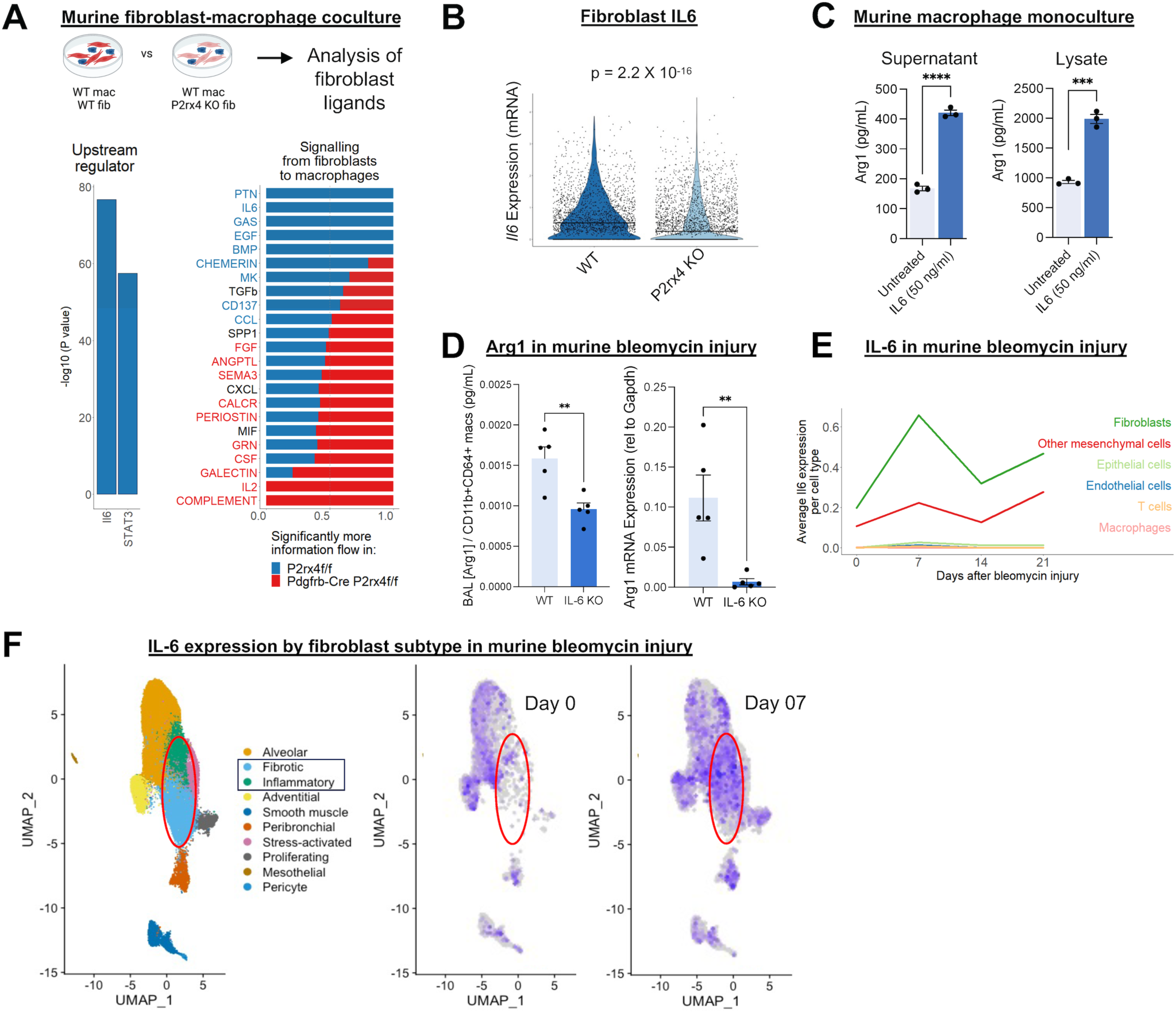
**IL-6 is necessary for Arginase 1 expression after bleomycin injury in mice.** A) Left: Ingenuity Pathways Analysis of predicted upstream regulators for macrophages cultured with WT relative to P2rx4 KO fibroblasts. Right: CellChat plot comparing WT and P2rx4 conditions. Significance was determined by Wilcoxon test with p<0.05. Data represent n=2 separate cocultures. B) Violin plot for IL-6 expression in WT relative to P2rx4 KO fibroblasts. Line shows median values. Data represent n=2 separate cocultures. C) Arg1 ELISA of cell lysates and conditioned media collected from mouse lung macrophage monoculture 72 hours after IL-6 treatment. N=3 biological replicates per condition. ***p<0.001, ****p<0.0001 by Student’s t-test. +/- SEM. D) Left: Arg1 ELISA of bronchoalveolar lavage fluid normalized to lung CD11b+CD64+ macrophage count taken at day 14 from bleomycin-injured WT or IL6 KO mice. Right: Arg1 qPCR of CD11b+CD64+ macrophages. N=5 mice per condition. **p<0.01 by Student’s t test. +/- SEM. E) Quantification of *Il6* expression by cell type in cells sequenced by scRNAseq directly after isolation from the lung at steady state and multiple time points after injury (reanalysis of merged data(8, 9) normalized by Sctransform(62)). F) UMAP of fibroblast subtypes (left) and IL-6 feature plots at steady state (center) and 7 days after bleomycin injury (right), from Tsukui et al.(8)

To test whether IL-6 could induce ARG1 in human lung, we first tested PCLS from explanted IPF lungs. Treatment with the IL-6R blocking antibody tocilizumab for 24 hours decreased ARG1 expression in IPF PCLS lysates (**Figure 5A**). To confirm that paracrine interaction between ARG1+ and IL-6+ cells was spatially plausible, we analyzed spatial transcriptomic data of IPF samples. For this purpose, we defined a proximity statistic that allowed us to quantitatively compare the probabilities of detecting an IL6+ fibroblast versus an IL6-fibroblast, within a given distance from an ARG1+ cell (**Methods**). Interestingly, at a distance from ARG1+ cells consistent with potential paracrine signaling (25 microns), fibroblasts were significantly more likely to express IL-6 than not to express IL-6 (**Figure 5B**). These data are consistent with paracrine IL-6 signaling from fibroblasts to macrophages resulting in ARG1 induction in IPF.

**Figure 5:**
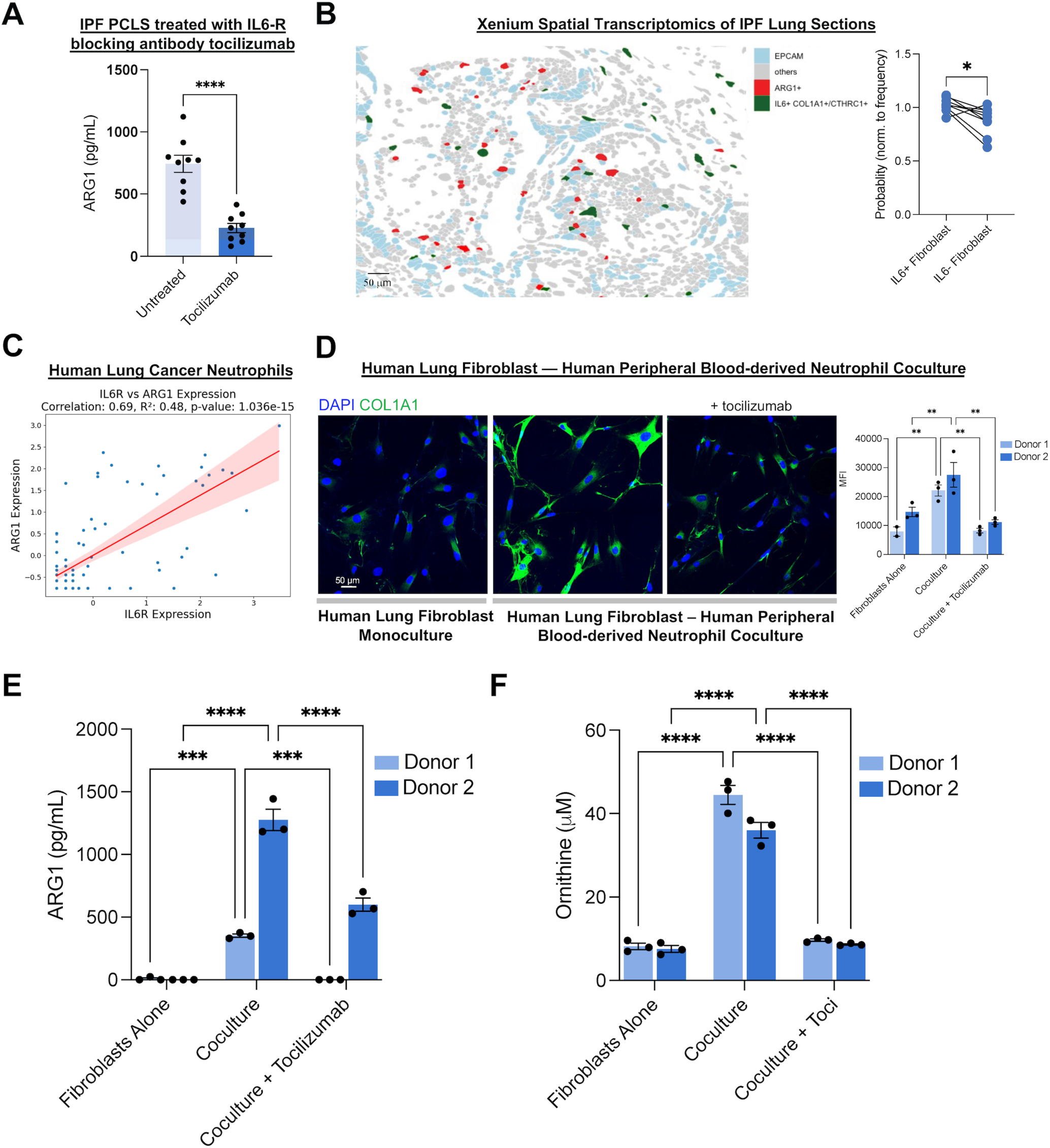
**IL-6 is necessary for Arginase 1 expression in IPF lung and in cocultures of human blood-derived neutrophils and lung fibroblasts.** A) ARG1 ELISA of lysates of precision-cut IPF lung slices cultured for 24 hours with or without IL-6R blocking antibody tocilizumab. Data represent N=3 patients, with 3 slices per condition per patient assayed. ****p<0.0001 by Student’s t-test. +/- SEM. B) 10x Xenium spatial analysis: Left: Representative field of view showing cells expressing ARG1, EPCAM, and IL6+ fibroblasts (defined by co-expression of either COL1A1 or CTHRC1). Right: Proximity analysis of Xenium data: Data shown are for the probability ratio, P (exp | Arg1+)/P(exp), the probability of encountering an IL6+ fibroblast, normalized by fibroblast frequency, at the radial distance of 25 μm from Arg1+ cells. *p<0.05 by paired Student’s t-test. Data are for N=9 separate patients. +/- SEM. C) Correlation scatter plot of tissue-resident neutrophils expressing IL-6R and ARG1. Represents integrated scRNAseq data of non-small cell lung cancer (NSCLC) from 19 datasets (32). D) COL1A1 immunofluorescence of human lung fibroblasts that were either monocultured, or cocultured with neutrophils with or without IL-6R blocking antibody (tocilizumab) treatment. Quantification is for n=3 separate cultures each. Donor 1 and Donor 2 represent separate donors for both neutrophils and fibroblasts.**p<0.01 by 2-way ANOVA followed by Sidak’s multiple comparisons test. +/- SEM. E) ARG1 ELISA of conditioned media from cocultures of human blood-derived neutrophils and fibroblasts from healthy human explanted lung. ***p<0.001, ****p<0.0001 by 2-way ANOVA followed by Sidak’s multiple comparisons test. +/- SEM. F) Ornithine concentration of conditioned media from the same cocultures as in (E). ****p<0.0001 by 2-way ANOVA followed by Sidak’s multiple comparisons test. +/- SEM.

We then interrogated published datasets for neutrophils from lung tissue to test whether ARG1+ neutrophils were more likely to express IL6R and therefore to support IL6 signaling. While genome-wide transcriptomic profiling of neutrophils from human fibrotic lungs has not yet been reported in the literature, as an alternative, we analyzed aggregate data from available non-small cell lung cancer (NSCLC) tumor microenvironment studies from 309 patients across 19 datasets (32). This analysis revealed a significant correlation between ARG1 and IL6R expression in neutrophils, consistent with the hypothesis that ARG1 expression in neutrophils is driven by IL-6 receptor ligation and signaling (**Figure 5C).**

Finally, we cocultured human peripheral blood-derived CD15+CD16+CD66B+CD14-neutrophils (**Supplemental Figure 3G**) with human lung fibroblasts. After 24 hours of coculture, we removed the neutrophils, which are nonadherent cells, and measured collagen expression in the fibroblasts by immunofluorescence. Notably, neutrophil coculture increased fibroblast collagen expression compared to fibroblast monoculture, and this increase could be blocked by IL6-R blockade with tocilizumab (**Figure 5D**). Notably, ARG1 and ornithine levels in the conditioned media were suppressed by tocilizumab (**Figure 5E-F**). Taken together, these analyses support the relevance of fibroblast IL-6-dependent ARG1 expression in the human lung fibrotic niche.

### eATP signaling induces fibroblast IL-6

To directly test whether the eATP receptor P2rx4 regulates fibroblast IL-6 expression, we considered that myeloid cells including both macrophages and neutrophils can be a source of eATP (33). Thus, we performed scRNAseq of fibroblasts cocultured with macrophages or cultured alone. Interestingly, coculture increased fibroblast expression of IL-6 (**Figure 6A**), and GO analysis confirmed that the GO pathway “Cellular Response to ATP” was enriched in fibroblasts in coculture compared to monoculture (**Figure 6B**). Importantly, IL-6 was higher in the conditioned media of WT macrophages cocultured with WT versus P2rx4 knockout (KO) fibroblasts (**Figure 6C**). Furthermore, direct treatment of cultured murine lung fibroblasts with ATPγS, a nonhydrolyzable form of ATP, increased IL-6 in the conditioned media of WT but not P2rx4 KO lung fibroblasts, and this effect was blocked by inhibition of p38 Map Kinase, which has been shown to mediate signaling downstream of P2rx4 (34, 35) (**Figure 6D**). To evaluate these results *in vivo*, we tested mice with fibroblast-specific P2rx4 knockout (KO) after bleomycin injury. We found reduced IL-6 levels in the bronchoalveolar lavage fluid of KO compared to WT mice (**Figure 6E)**. Furthermore, siRNA KD of P2RX4 in human lung fibroblasts decreased IL-6 expression in response to ATPγS (**Figure 6F)**. We also found that P2rx4 expression increased post-bleomycin injury (**Supplemental Figure 3H**), consistent with increased functional effects of this pathway during fibrogenesis. Finally, coculture of human peripheral blood-derived neutrophils with human lung fibroblasts increased IL-6 in the conditioned media compared to fibroblasts or neutrophils alone. This effect could be inhibited by treatment with a small molecule inhibitor of P2rx4 (BAY-1797(36); **Figure 6G**). Taken together, our findings indicate that, in both murine and human systems, eATP-P2rx4 signaling regulates fibroblast IL-6 expression, which in turn induces myeloid Arg1 resulting in ornithine loading of fibroblasts for proline synthesis and pathologic collagen expression.

**Figure 6:**
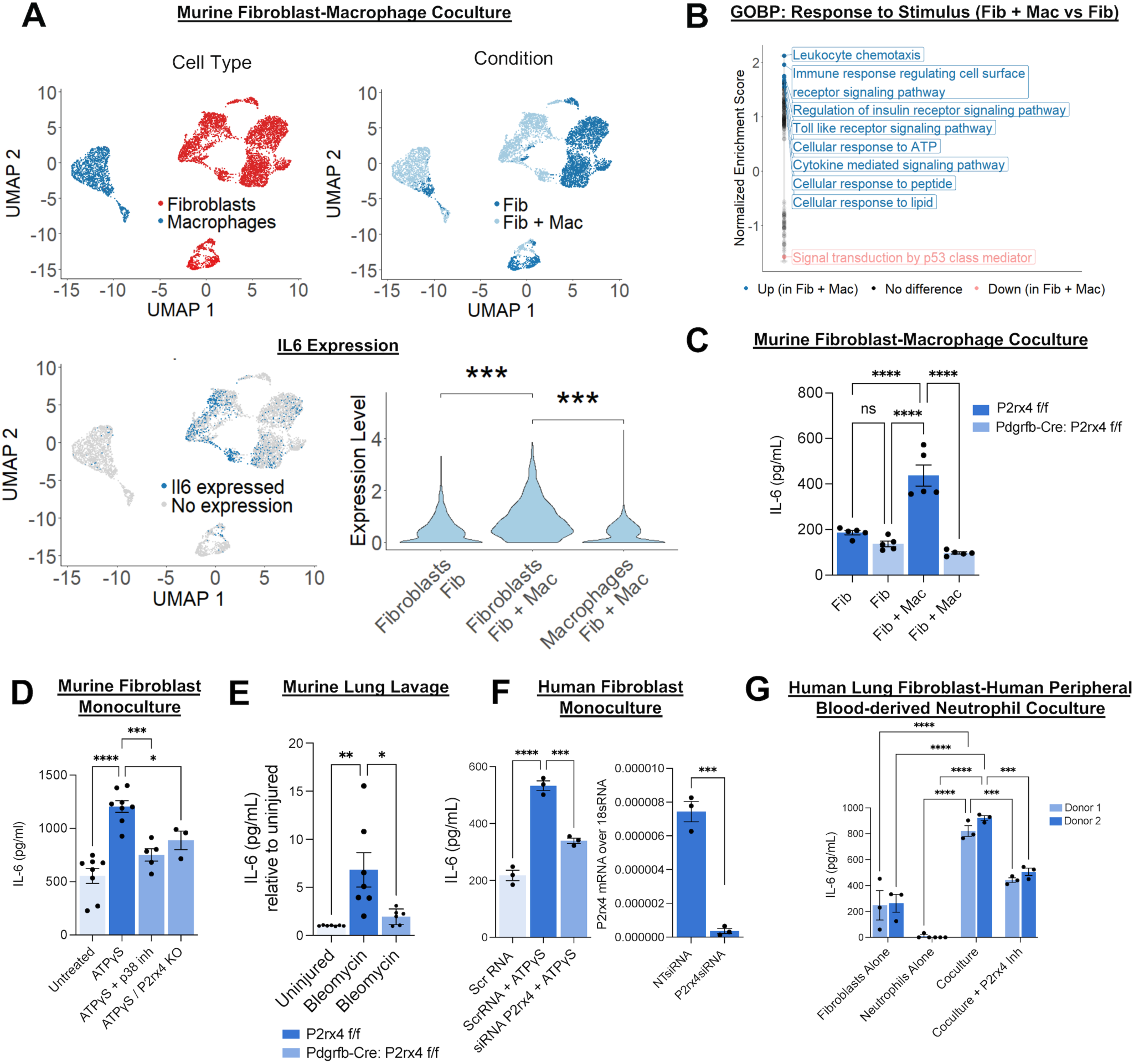
**P2rx4 is necessary for IL-6 expression in mouse and human lung fibroblasts.** A) Top Left: UMAP plot of scRNAseq for macrophage-fibroblast cocultures with SingleR-based cell type annotation(2) shown. Data represent n=2 separate cultures for each condition. Top Right: UMAP plot for data showing sample of origin (“Fib” = fibroblast monoculture, “Fib + Mac” = fibroblast coculture with macrophages). Below Left: Feature plot showing *Il6* expression. Below Right: Violin plot of *Il6 expression*. ***p<0.001 by Wilcoxon Rank Sum test corrected for multiple comparisons by Bonferroni method. B) Gene set enrichment analysis (GSEA) of fibroblast single cell transcriptomes in coculture with macrophages compared to fibroblast monoculture using GO “Response to stimulus” pathways. C) IL-6 ELISA of conditioned media from macrophage-fibroblast cocultures with or without fibroblast-specific P2rx4 deletion. N=5 five biological replicates per condition. ****p<0.0001 by 1-way ANOVA with post hoc Sidak’s multiple comparisons tests. +/- SEM. D) IL-6 ELISA of conditioned media from monocultured mouse lung fibroblasts from WT mice, with and without ATPγS and SB203580 (p38 Map Kinase inhibitor) treatment or from fibroblast-specific P2rx4 KO (Pdgfrb-Cre: P2rx4 f/f) mice, with ATPγS treatment. N=8, 8, 5, 3 biological replicates per condition. *p<0.05, ***p<0.001, ****p<0.0001 by 1-way ANOVA with post hoc Sidak’s multiple comparisons tests. +/- SEM. E) IL-6 ELISA from bronchoalveolar lavage from mice with or without fibroblast-specific P2rx4 deletion. N=7, 7, 6 biological replicates, left to right. **p<0.01, *p<0.05 by 1-way ANOVA with post hoc Sidak’s multiple comparisons tests. +/- SEM. F) IL-6 ELISA for conditioned media from human donor lung fibroblast monocultures with P2RX4 siRNA KD or scrRNA, +/-ATPγS treatment (*left*, n=3 per condition. ***p<0.001, ****p<0.0001 by 1-way ANOVA with post hoc Sidak’s multiple comparisons tests; quantification of KD by qPCR on *right*, n=3 per condition, ***p<0.001 by Student’s t-test). +/-SEM. G) IL-6 ELISA for conditioned media from human donor lung fibroblast monocultures, human blood derived neutrophil monocultures, or cocultures with or without P2RX4 inhibition with BAY-1797. Conditioned media were collected after 24 hours of culture. Each point re[presents a separate technical replicate. Neutrophils were derived from 2 separate blood donors, and fibroblasts were derived from 2 separate healthy lung donors. ***p<0.001, ****p<0.0001 by 2-way ANOVA followed by Sidak’s multiple comparisons test. Donor 1 and Donor 2 represent separate donors for both neutrophils and fibroblasts. +/- SEM.

## Discussion

For the first time to our knowledge, we demonstrate that Arg1 and its enzymatic product ornithine have a profibrotic effect by driving fibroblast proline synthesis, a key substrate for collagen expression.

Imaging studies indicated that ARG1 is expressed more highly in IPF than in healthy human lung explants. Studies using both genetic deletion and chemical inhibition demonstrated the profibrotic effect of Arg1 in murine lung injury and in IPF precision-cut lung slices. Remarkably, we found that the reaction product of Arg1, ornithine, was increased in fibrotic lung tissue from mice and IPF patients.

These data build on the recent findings that ornithine was the only amino acid to be elevated in plasma of 100 IPF patients compared to 300 healthy controls (37) and that the expression of ornithine aminotransferase, which converts ornithine to the proline precursor P5C, was correlated with areas of fibrosis in IPF lung (38). Furthermore, we found that ornithine had a profibrotic effect: Ornithine directly increased collagen expression *in vivo* and in cultured murine and human lung fibroblasts. Metabolic labeling studies revealed that ornithine was a substrate for proline synthesis.

Our findings also address how this profibrotic amino acid metabolism was initiated via paracrine crosstalk. In both murine and human fibrotic lung and coculture systems, myeloid Arg1 expression was dependent on IL-6 expression in fibroblasts, which in turn was regulated by eATP-P2rx4 signaling. The recent literature has revealed a subset of fibroblasts expressing inflammatory genes in fibrosis, including cytokines (7, 8). A pathologic function of fibroblast-derived cytokine expression is elucidated by our discovery that fibroblast IL-6 is necessary for Arg1 expression in myeloid cells, with the profibrotic consequence of ornithine loading of fibroblasts leading to increased collagen production.

The role of type 2 inflammation in fibrosis deserves mention with respect to Arg1. In schistosomiasis-induced fibrosis models, where Arg1 deletion led to an increase in type 2 inflammation and consequently increased fibrosis because of an accumulation of arginine, which supports CD4 T cell proliferation (39). However, in IPF and in murine bleomycin-induced pulmonary fibrosis, type 2 inflammation does not appear to be an important driver of fibrosis: Inhibition of type 2 inflammation with dual IL-4 and IL-13 blockade failed to show efficacy in a clinical trial of IPF (40). Furthermore, in the bleomycin-induced sterile injury model that we have used, the onset of type 2 inflammation is later than the period when significant collagen deposition has already occurred and is therefore less relevant to fibrogenesis (41). Therefore, we conclude that type 2 inflammation may be less important in the model.

The myeloid cell type expressing Arg1 varied by species. In mice, we found that Arg1 was predominantly expressed in CD11b+CD64+ macrophages and not in canonical Ly6G+ neutrophils. However, there was Arg1 expression in a relatively smaller population of Ly6G+CD64+CD11c+ cells, which may be an intermediate inflammatory myeloid lineage similar to a profile that others have recently described in the setting of murine influenza infection (42). Future studies should more deeply profile these latter cells and ARG1+ cells from IPF, which expressed neutrophil markers, to determine whether they are phenotypically similar. Nonetheless, in both species we confirmed a P2RX4 – IL-6 – ARG1, myeloid-mesenchymal circuit that is functionally important in lung fibrosis. Notably, blood and bronchoalveolar lavage neutrophilia has been associated with fibrosis progression and worse prognosis in IPF (20–23). Our work highlights ARG1 and ornithine metabolism as a neutrophil-dependent pathway that should be developed as a potential therapeutic target for IPF.

There are limitations that should be considered when interpreting our results. First, our metabolomic studies indicate conversion of ornithine to proline, a key substrate for collagen synthesis; however, we note that ornithine conversion to polyamines with profibrotic potential remains a further possibility not yet addressed by our results. Second, Arg2 is another arginase that could also contribute to ornithine loading of fibroblasts, and future studies should focus on how both arginases are regulated across the temporal phases of fibrosis. Third, our findings are agnostic with respect to the site of ornithine generation; we detected Arg1 both in both lysates and conditioned media of IL-6-treated macrophages, consistent with the possibility of both intracellular ornithine production and export as well as extracellular production, as per published reports (43, 44). Finally, regarding sources of eATP, we concede that it is likely that multiple sources of eATP exist in the fibrotic niche, such as dying or dysfunctional cells, in addition to myeloid cells themselves. Our results nonetheless reveal that the DAMP signal eATP triggers expression of fibroblast IL-6, which induces profibrotic Arg1 in neighboring myeloid cells.

In conclusion, we show the dependence of lung fibrosis on ornithine deriving from myeloid Arg1 in mouse models and in functional studies of IPF lung. The importance of understanding the contribution of amino acid metabolism to fibrosis has been recognized in recent years (45), and our findings highlight the role of its immune regulation in the synthesis of pathologic collagen. These studies increase enthusiasm for targeting ARG1 as a therapeutic approach in IPF.

## Methods

### Sex as a biological variable

For preclinical in vivo studies, we used both female and male mice, and experimental groups were balanced with respect to sex. Human lung samples were acquired from both males and females.

### Human lung tissues

IPF lung samples were obtained as explants at the time of lung transplantation. Deceased donor-control lungs not known to have lung disease were made available by Donor Network West. Demographic data with respect to ethnicity and race (**Supplemental Table 1**) were derived from the electronic medical record and classified as per NIH notice NOT-OD-15-089.

### Mice

Pdgfrb-Cre(46), Lysm-Cre (47), P2rx4 floxed (48), Arg1 floxed (49), Col1a1-EGFP (50), Arg1-RFP-CreERT2 (51), and Arg1-YFP (52) mice were available to the investigative team, and IL-6 KO (53), WT, and R26-LSLS-TdTomato mice were obtained from JAX. Col1a1-GFP mice were obtained from David Brenner (50). All mice were maintained on a C57BL/6 background and were maintained in specific pathogen-free animal barrier facility at the University of California, San Francisco. All experiments were performed on 6-8 weeks old, sex-matched mice.

### Mouse lung injury

For lung injury, mice anesthetized with isoflurane were instilled intratracheally with bleomycin (Fresenius, 3U/kg). In the case of Arg1 inhibition, bleomycin-injured mice were treated daily from day 9 to day 15 post-injury with 100 mg/kg CB-1158 (Numidargistat dihydrochloride, HY-101979A, MedChemExpress) dissolved in water, by oral gavage. In the case of ornithine treatment, mice were treated twice daily by gavage with ornithine 2 g/kg dissolved in 100 mL of water.

### Murine macrophage-fibroblast coculture

We prepared cocultures of macrophages and fibroblasts that were isolated from the lungs of mice 7 days after bleomycin lung injury. To make single cell suspensions, lungs were harvested from euthanized mice and minced with scissors, and the tissue was resuspended in RPMI with 0.2% Collagenase (10103586001, Roche), 2000 U/ml DNase I (4716728001, Roche), and 0.1 mg/ml Dispase II (4942078001, Sigma). The suspension was incubated for 1 hour at 37^ο^C and then passed through a 70 μm filter (130-110-916, MACS SmartStrainers) to obtain single cells. The cells were washed twice with 1X PBS pH 7.4 (10010023, Gibco). For macrophage isolation, a total of 10^7^ cells were resuspended in 80μl PBS buffer supplemented with 0.5% BSA (0332, VWR Life Science) and 2mM EDTA (E57040, Research Products International). 20μl CD11b microbeads were used per 10^7^ cells. Macrophages were obtained by positive selection by binding to CD11b microbeads and passing through MACS columns (130-049-601, Miltenyi Biotech).

Isolated CD11b^+^ macrophages were cultured in DMEM supplemented with 10% FBS, 1% penicillin-streptomycin (GIBCO), and 20 ng/ml M-CSF (315-02, Peprotech) for 2 days. Primary mouse lung fibroblasts were freshly isolated after magnetic bead-based negative selection of epithelial cells (118204, Biotin anti-mouse CD326 Epcam, Clone: G8.8, BioLegend), endothelial cells (102404, Biotin anti-mouse CD31, clone: 390), immune cells (103104, Biotin anti-mouse CD45, clone: 30-F11, BioLegend), pericytes and smooth muscle cells (134716, Biotin anti-mouse CD146, clone: ME-9F1, BioLegend), and red blood cells (116204, Biotin anti-mouse Ter119, clone: TER-119, BioLegend) with biotinylated antibodies and Dynabeads (MyOne Streptavidin T1, 65601, Thermo Fisher Scientific) as previously described (7). Fibroblasts were cultured for 5 days prior to trypsinization and addition to macrophages for coculture at a 1:1 ratio in DMEM complete media for 5 further days.

### Human lung fibroblast isolation

Human lung tissue from deceased donors not used for transplant was resected and minced in Hanks’ Balanced Salt Solution (HBSS) buffer supplemented with 0.2% Collagenase (10103586001, Roche), 2000 U/ml DNase I (4716728001, Roche), 0.1 mg/ml Dispase II (4942078001, Sigma), and 1% Penicillin-streptomycin for 1.5 hours at 37^ο^C and 5% CO2. 1X Amphotericin B (15290026, Gibco) was added to the dissociation solution for the last 30 min. Digested lung tissue was lysed further with a MACS tissue dissociator (gentleMACS Dissociator, Miltenyi Biotec) using gentleMACS C tubes (130-093-237, Miltenyi Biotec) at mLUNG-01 setting. The suspension was then passed through a 70 μm filter to obtain single cells. Cells were resuspended in PBS with 0.5% BSA and 2mM EDTA. For the negative selection of human fibroblasts, the following antibodies were used at a 1:200 ratio: epithelial cells (324216, Biotin anti-human Epcam, clone:9C4, BioLegend), endothelial cells (13-0319-82, Biotin anti-human CD31, clone: WM-59, Invitrogen), immune cells (368534, Biotin anti-human CD45, clone: 2D1, BioLegend), and pericytes and smooth muscle cells (361036, Biotin anti-human CD146, clone: P1H12, BioLegend). Fibroblasts were cultured in DMEM supplemented with 10% serum, 1% penicillin-streptomycin, and 1% amphotericin B for a total of 5 days.

### Human neutrophil isolation and coculture with fibroblasts

Whole blood (10ml) from heathy donors was collected in BD Vacutainer® k-EDTA Tubes (Vitalant, San Francisco, CA) and used for primary neutrophil isolation as previously described (54). Briefly, 7ml blood was layered on top of 7ml of PolymorphPrep (Cosmo Bio Usa Inc, AXS-1114683) and centrifuged at 500 x *g* for 35 min at room temperature without braking. The peripheral blood mononuclear cells and plasma layers were aspirated and the neutrophil layer was collected. The cells were washed with PBS and centrifuged at 400g for 5 minutes. The cell pellet was resuspended in 3ml ACK lysis buffer (Thermo Fisher, A1049201) and centrifuged at 400g for 5 minutes. Neutrophil purity was 95%, confirmed by flow cytometry (**Supplemental Figure 3F;** Antibodies: CD15 clone W6D3, Biolegend cat# 323039; CD66B clone G10F5, Biolegend cat# B221034; CD16 clone 3G8, BD Biosciences cat# 560474; CD14 Clone 63D3, Biolegend cat# 367118). The cells were finally resuspended in RPMI supplemented with 10% FBS and counted. Neutrophils were then added to human lung fibroblasts for 24 hours of coculture in RPMI with 10% FBS, with or without P2RX4 inhibitor BAY-1797 (1 μm, 7573, Tocris) or IL6-R blocking antibody tocilizumab (100 ng/mL, HY-P9917, MedChemExpress). The conditioned media were retained for analysis after separation of the cellular fraction by centrifugation, and fibroblasts were fixed and permeabilized for immunofluorescence analysis.

### Flow cytometry and sorting

*Cocultured cells:* Cocultures were trypsinized with 0.25% trypsin and resuspended in 1X PBS with 0.5% BSA and 2mM EDTA. Single-cell suspensions were pre-stained with Fc blocker for 10 min in ice followed by staining at 1:300 with anti-CD64 (139304, PE anti-mouse CD64, clone:X54-5/7.1, BioLegend) and anti-Pdgfra (25-1401-82, PE-Cy7, anti-mouse CD140a, clone: APA5, Invitrogen) antibodies for 40 min. DAPI was used to distinguish dead cells. *Mouse lung cell suspensions:* Whole lung single cell suspensions were prepared by harvesting lung lobes into 5 ml HBSS with 40µl Liberase Tm (0.1 U/ml, 5401127001, Roche) and 20µl DNAse 1 (10mg/ml, Roche, Cat# 10104159001), followed by automated tissue dissociation (GentleMacs; Miltenyi Biotec) and tissue digestion for 30 min at 37°C on a shaker. Digested samples were processed on the GentleMacs using the “lung2” program, passed through 70µm filters, and washed, followed by red blood cell lysis and final suspension in FACS buffer. Cells were counted using a NucleoCounter (Chemometic). All samples were stained in 96-well V-bottom plates. Single cell samples were first incubated with antibodies to surface antigens for 30 min at 4°C in 50µl staining volume. Flow cytometry was performed on BD LSRFortessa X-20. Fluorochrome compensation was performed with single-stained UltraComp eBeads (Invitrogen, Cat# 01-2222-42). Samples were FSC-A/SSC-A gated to exclude debris, followed by FSC-H/FSC-A gating to select single cells and Draq7 viability dye (Biolegend) to exclude dead cells. Monocyte-derived macrophages (moMacs) were identified as CD19^-^, Ly6G^-^, NK1.1^-^, Siglec-F^-^, CD11b^+^, and CD64^+^. Arginase1-positive cells were identified by the presence of eYFP. Data were analyzed using FlowJo software (TreeStar, USA) and compiled using Prism (GraphPad Software). Monoclonal antibodies used were: anti-CD45 (30-F11, Biolegend), anti-CD11b (M1/70, Biolegend), anti-CD11c (N418, Biolegend), anti-NK1.1 (PK136, Biolegend), anti-CD19 (6D5, Biolegend), anti-CD64 (X54-5/7, eBiosciences), anti-Ly6G (1A8, Biolegend), anti-SiglecF (E50-2440), anti-I-A/I-E (MHCII) (M5/114.15.2, Biolegend), and anti-CCR2 (475301, BD BioSciences). In the case of quantifying and sorting CD45+CD11b+CD64+ monocyte-derived macrophages in bleomycin-injured WT and IL-6 KO mouse lung cell suspensions, 50,000 microbeads (Invitrogen) were added to each sample for quantification of absolute live CD45+CD11b+CD64+ cells. Lavage cell pellets were added to the suspensions prior to sorting. Cells were sorted on a FACS Aria2.

### Quantitative Real-Time PCR analysis

Human lung fibroblasts were lysed in 300μl TRIzol reagent (10296010, Ambion life technologies) to obtained RNA. 600 ng of RNA per sample was used to prepare cDNA using iScript reverse transcriptase supermix (Bio-Rad, 1708841). qPCR was performed for the target genes using SYBR Green super mix (Applied Biosystems, 4309155). Gene expression was normalized to 18S rRNA. The following primers were used: P2RX4 Forward: GAGATTCCAGATGCGACCACT P2RX4 Reverse: ACCCGTTGAAAGCTACGCAC 18S rRNA Forward: GTAACCCGTTGAACCCCATT 18S rRNA Reverse: CCATCCAATCGGTAGTAGCG In the case of sorted CD45+CD11b+CD64+ murine lung monocyte-derived macrophages, mRNA was isolated form pelleted cells as above, and qPCR was performed. Gene expression was normalized to Gapdh. The following primers were used:

Arg1 Forward: CTCCAAGCCAAAGTCCTTAGAG Arg1 Reverse : AGGAGCTGTCATTAGGGACATC Gapdh Forward: AGTATGACTCCACTCACGGCAA Gapdh Reverse: TCTCGCTCCTGGAAGATGGT

### siRNA Knockdown

Healthy human fibroblasts were treated with 5μM siRNA (Dharmacon, D-001810-01-05 ON-TARGET plus No-targeting siRNA) control or (Dharmacon, L-006285-00-0010 ON-TARGET plus Human P2RX4 SMARTpool) using DharmaFECT in serum-free media for 12 hours, followed by supplementing with DMEM with 10% serum. Cells were incubated for an additional 48 hours. Treated cells were harvested for either RNA isolated or stimulated with 100μM ATPγS for 12 hours and culture supernatant was collected for Human IL6 ELISA assay.

#### IL-6 ELISA

Bronchoalveolar lavage fluid from mice or conditioned media from cultured murine cells were collected for measurement of IL-6 with a Duoset ELISA kit (R&D Systems, DY406-05). Conditioned media from cultured human cells were collected and IL-6 was measured using Duoset ELISA kit (R&D Systems, DT206-05). In both cases, the capture antibody was coated overnight in a 96-well plate at room temperature. Wells were aspirated and washed in ELISA wash buffer and blocked by adding reagent diluent consisting of 1% BSA in PBS. After 1 hour of room temperature incubation, wells were again washed with wash buffer and samples or standards were added and incubated for 2 hours at room temperature. Detection antibody was used along with streptavidin-HRP and substrate solution. The reaction was stopped with stop solution, and optical density was measured using a microplate reader at 450 nm.

### Arginase 1 ELISA

*Mouse:* Arginase 1 levels were measured from conditioned media or from bronchoalveolar lavage, using Mouse Arginase 1 ELISA Kit (ab269541, Abcam) based on the manufacturer’s protocol. Briefly, CD11b+ mouse lung cells were seeded in 12-well dishes for 2 days in DMEM supplemented with M-Csf (315-02, Peprotech). Cells were exposed to murine IL-6 for 72 hours, and culture supernatant and cell lysate were then collected. For *in vivo* experiments, bleomycin-injured mice received bronchoalveolar lavage with 1 mL of PBS on day 14 after injury, and the noncellular fraction was retained for analysis. For the ELISA, subsequently, 50 μl of undiluted samples were added to each well, along with a 50 μl antibody cocktail comprising capture antibody and detection antibody dissolved in antibody diluent. Plates were incubated for 1 hour at room temperature. Wells were washed thrice with wash buffer and incubated with 100 μl Tetramethylbenzidine (TMB) substrate for 15 min. The reaction was stopped using 100μl stop solution, and OD was measured at 450nm. *For Human:* Arginase 1 levels were measured from IPF precision-cut lung slice (PCLS) using the Human Arginase 1 ELISA Kit (BMS2216, ThermoFisher), following the manufacturer’s instructions. Briefly, following 24 hours of culture with or without tocilizumab (100 ng/mL, HY-P9917, MedChemExpress), PCLS were lysed in RIPA buffer with protease inhibitor, and 100 μl of standards or undiluted sample was added to each well of a pre-coated 96-well plate and incubated for 2 hours at room temperature.

Wells were then washed four times with provided wash buffer before the addition of 100 μl of biotin-conjugated anti-arginase 1 antibody to each well, followed by a 1-hour incubation. After another wash step, 100 μl of streptavidin-horseradish peroxidase (HRP) was added and incubated for 30 minutes. Wells were washed again and incubated with 100 μl of TMB substrate solution for 15 minutes. The enzymatic reaction was stopped by adding 100 μl of stop solution, and optical density (OD) was measured at 450 nm using a microplate reader.

### Histology and Immunofluorescence imaging

*Mouse lung:* Arg1-RFP-creERT2 /Rosa26-LSL-TdTomato/Col1a1-EGFP mice were injured for 14 days. Lungs from uninjured or injured mice were inflated with a solution consisting of 30% sucrose solution mixed 1:1 with OCT (4585, OCT) and fixed in 4% formaldehyde at 4^ο^C for 4 hours. Lungs were then washed and submerged in 30% sucrose solution overnight at 4^ο^C. Next, the tissue was incubated in the 1:1 solution of 30% sucrose and OCT overnight followed by changing to OCT for 2 hours, and then a tissue block was frozen after embedding in OCT. 5 μm sections were cut from OCT-embedded tissue. For immunofluorescence imaging, frozen sections were incubated with Mertk antibody (AF591, R&D Systems) or S100A8 antibody (AF3059, R&D Systems) in staining buffer (1% BSA, 0.5% Triton x-100 in PBS), washed, and then incubated with anti-rabbit secondary antibody (Invitrogen). Slides were washed with PBS and mounted on antifade DAPI mounting media. *Human lung:* 5 μm thick sections from patient FFPE blocks were collected for IHC staining. Slides were stained using Opal Manual IHC kit (PerkinElmer). After deparaffinization, antigen retrieval was performed in AR buffer for 45 sec at 100% power followed by 15 min at 20% power. After blocking slides were incubated with primary antibodies ARG1 (Cat# 93668, Cell Signaling Technology) overnight at 4C. Polymer HRP was introduced to slides for 10 min, followed by signal amplification using Opal570 (NEL810001KT,Akoya Biosciences) for 10 min at room temperature. The slides were then counterstained with DAPI mounting media and scanned using with a 10x and 40x objective of Leica inverted widefield microscope. *Cultured Cells:* Macrophages and fibroblasts or fibroblasts alone, isolated from mouse lung as detailed above, were cultured on glass coverslips and in some cases treated with CB-1158 (1 μM), ornithine (1 mM), or 1 μM OAT inhibitor 5-FMOrn dihydrochloride (HY154021A, MedChemExpress). Cells were fixed with 4% PFA, permeabilized with 1% BSA, 0.5 % triton x-100 in PBS, and incubated with primary antibodies (Col1a1, #720265, Cell Signaling Technology; α-SMA, A5228, Sigma) followed wash, secondary antibodies, further wash, and mounting. Imaging was performed on a Leica SP8 laser scanning confocal microscope. Fluorescence signal was quantified with Imaris software for cellular areas.

### Western blot

Precision cut lung slices were obtained from human IPF tissue sections. Briefly, IPF tissue was injected with 2% low melting agarose (50111, Lonza) and then submerged in ice-cold PBS to allow solidification of agarose. 400 μm lung slices was generated using a Compresstome device (VF-310-OZ, Precisionary Instruments). Slices were kept in complete DMEM media for 2 hours to allow the release of agarose followed by changing to fresh complete DMEM with 10% FBS and 1% penicillin-streptomycin. Lung slices were treated with 50 μM CB-1158 for 24 hours. After incubation slices were minced using a tissue homogenizer. Cells were lysed in Pierce RIPA buffer (Thermo Fisher Scientific, 89901) with protease inhibitor cocktail (Thermo Fisher Scientific, 1861278). 10 μg of protein was run on 10% sodium dodecyl sulfate-polyacrylamide gel electrophoresis (Bio-rad, 4561034) and transferred to a PVDF membrane (Thermo Fisher Scientific, 88520). The membrane was incubated with 1:1000 Col1a1 antibody (#720265, Cell Signaling Technology) overnight at 4^ο^C. Blots were washed in 1X TBST and incubated with peroxidase-conjugated goat anti-rabbit (1:20,000, Anaspec, AS28177) for 4 hr at 4^ο^C. These blots were developed using SuperSignal West Pico Chemiluminescent substrate (Thermo Fisher Scientific, 34080) in ChemiDoc XRS+ gel imaging system (Bio-rad). Quantification of bands was done using ImageJ.

### Hydroxyproline assay

Mice were euthanized and lungs were excised and snap-frozen. Isolated lung samples were homogenized and incubated with 50% trichloroacetic acid (T6399, Sigma) on ice for 20 min. Samples were then incubated in 12N HCL (A144, Fisher) overnight at 110^ο^C. Dried pellets were reconstituted in distilled water with constant shaking for 2 hours at room temperature. Samples were then mixed with 1.4% Chloramine T (Sigma, 85739) and 0.5 M sodium acetate (Sigma, 241245) in 10% 2-propanol (Fisher, A416) and incubated with Ehrlich’s solution (Sigma, 03891) for 15 min at 65°C. Absorbance was quantified at 550 nm, and concentration was calculated using a standard curve of commercial hydroxyproline (Sigma, H5534).

### scRNAseq library preparation and sequencing

PIPseq T2 3ʹ Single Cell RNA Kit (v3.0 – cultured cells, v4.0 – directly sequenced cells) was used for pre-templated instant partitioning (PIP) to capture single-cell mRNA transcripts with PIP beads according to manufacturer’s protocol (Fluent Biosciences, FB0001026). After 5 days of coculture, single cell suspension of cells was obtained by trypsinization followed by washing in 1X PBS. Cells were washed in ice-cold PIPseq Cell Suspension Buffer (Fluent Biosciences, FB0002440). Cells were counted and stained with Trypan blue to confirm >90% viability. Single-cell library preparation was performed using the manufacturer-recommended default protocol and settings. The sequencing libraries were submitted to UCSF Center for Advanced Technology (Novaseq X) or Novogene (Novaseq 6000) for sequencing. The demultiplexed FASTQ files were aligned to mouse genome (mm10) using PIPseeker 1.0.0 (Fluent Biosciences). After sequence alignment, we observed around 50% of input cells being called when PIPseeker sensitivity level was set to 3, which was near the inflection point of the barcode rank plot. The original FASTQ files, the quality reports, and the expression matrix outputs of PIPseeker have been deposited in Gene Expression Omnibus (GEO).

### scRNAseq Data Analysis

The Seurat – Guided Clustering Tutorial (March 27, 2023) was followed to convert our expression matrixes into Seurat objects (Seurat version 4.3.0) (55). For quality control, we removed the cells with less than 200 genes or more than 10,000 genes and larger than 5% mitochondrial content. Seurat objects were integrated following Seurat’s Introduction to scRNAseq Integration (March 27, 2023), selecting the top 2,000 variable features as integration anchors. Cell doublets were removed with the package DoubletFinder (2.0.3) (56). Following Seurat – Guided Clustering Tutorial (March 27, 2023), we selected the top 2000 highly variable genes to obtain the cell UMAP coordinates and group the cells into clusters with a sensitivity of 0.5(55). Cell types were annotated using the SingleR package 1.10.04(2) using the ImmGen database (57) as reference. The Gene Set Enrichment Analysis (GSEA) was performed using the package enrichR 3.1(58) (FDR<0.05), with gene ontology data taken from the database “GO_Biological_Process_2021”(59). Cell communication pathways analysis was performed using the package CellChat (1.6.1) (27). Upstream regulator prediction was done using the Ingenuity Pathway Analysis software (Qiagen) (60). Differentially expressed genes for macrophages with p < 0.05 and average log2 fold change > 0.75 were used for analysis. Analysis results with p < 0.05 were considered significant.

### Multiplexed Ion Beam Imaging and Analysis (MIBI)

*Slide Preparation:* Serial 5 um FFPE sections were cut onto one glass and one gold slide. Both slides were baked at 70* Celsius overnight and deparaffinized in three washes of fresh Xylene and rehydrated in EtOH (2X 100% EtOH, 2X 90% EtOH, 1X 80% EtOH, 1X 70% EtOH) and distilled water (2X) washes. Washes were performed in (include wash machine specs). An antigen retrieval slide chamber was prepared by diluting 10X Tris with EDTA antigen retrieval buffer 1:10 in diH2O. The prepared slide chamber was added to a PT Module (add specs) filled with PBS and preheated to 75°C. The rehydrated slides were run in the preheated PT module at 97°C for 40 minutes, then cooled to 65°C in the PT module. The prepared slide chamber was then removed from the PT module and cooled at RT for 30 minutes. Slides were washed 2X in 1X TBS-Tween. *Glass Slide IHC:* Tissues were encircled by PAP pen boarders and blocked with 5% donkey serum diluted in TBS IHC wash buffer for 1 hour at RT in a moisture chamber. Wash buffer was aspirated from the slide, and Arginase-1 primary antibody (Ionpath Cat#: 715001) was stained overnight at 4°C overnight in the moisture chamber. The following day, primary antibody was aspirated, and slides were washed twice with 1X TBS-T for 5 minutes, blocked with 3% peroxide buffer for 15 minutes at RT, and washed again twice with 1X TBS-T for 5 minutes each wash before incubation with anti-rabbit secondary antibody for 1 hour at RT. Slides were washed twice with 1X TBS-T for 5 minutes before visualization with 100uL DAB for 5 minutes at RT. DAB reaction was stopped by tapping waste into contained waste bin, and then washing the slide into a slide chamber filled with diH2O 3X for 30 seconds each wash. The slide was then stained with Hematoxylin for 1 minute at RT and washed with tap water 2X for 30 seconds. The slide was then dehydrated by washing in EtOH (1X 70% EtOH, 1X 80% EtOH, 2X 95% EtOH, 2X 100% EtOH), and Xylene (2X) before cover slipping. *Gold Slide Staining:* Gold slides were transferred the Sequenza Immunostaining Center Staining System (Electron Microscopy Sciences, Hatfield, PA). Endogenous biotin-binding proteins with blocked with (Avidin/biotin-blocking reagents) for 30 minutes at room temperature. 5% donkey serum was added to the top of the chamber to wash out the avidin blocking reagents and block additional non-specific antibody binding sites for 1 hour at RT. Primary and Secondary antibodies panels were assembled with appropriate volumes of each titrated antibody and a final concentration of 0.05M EDTA. The complete cocktails were filtered through a pre-wet 0.1 um Ultragree MC Spin Filter, and then the Primary Antibody cocktail (**Supplemental Table 2**) was added to the Sequenza top chamber and incubated overnight at 4°C. The Secondary antibody cocktail was stored at 4°C. The following day, the slides were washed by adding 1X TBS-T to the Sequenza chamber 2X before adding the Secondary Antibody cocktail to the Sequenza chamber for 1 hour at RT. Gold slides were removed from the Sequenza chamber and washed 3X with 1X TBS-T for 5 minutes each wash, 1X with filtered 2% Glutaraldehyde for 5 minutes, 3X with filtered 1X Tris pH 8.5, 2X with filtered diH2O, 1X with 70% EtOH, 1X with 80% EtOH, 2X with 90% EtOH, and 2X with 100% EtOH. Slides were allowed to dry at RT for 10 minutes before being stored in a vacuum chamber (Cat#) before MIBIscope analysis. *Image Acquisition and Processing:* Gold slides were loaded into the MIBIscope (Ionpath, Menlo Park, CA) and FOVs were selected by matching tissue topography to ROIs with Arginase-1+ Staining from the serial IHC glass slide. FOVs were acquired at Fine resolution, with a dwell time of 1 second at a resolution of 0.39 um per pixel. Image QC was performed by following the Angelo Lab toffy pipeline (https://github.com/angelolab/toffy). Analysis was performed by following the Angelo Lab ark pipeline (https://github.com/angelolab/ark-analysis).

### Spatial Transcriptomics (10x Xenium)

FFPE preserved sections of lung tissue were prepared for spatial transcriptomics imaging on the Xenium platform by following 10x Genomics protocols CG000582 Rev E and CG000584 Rev A. Briefly, protocol CG000582 prepares the slides for the imaging run by first hybridizing the probe panel of choice. Here, the standard human lung panel available from 10x Genomics (Cat #1000601) was supplemented with a custom panel specific for lung disease states, including pulmonary fibrosis (**Supplemental Table 3**). Following probe hybridization, annealed probes are ligated together to create circular fragments, then circularized probes are amplified with a rolling circle PCR. After rolling circle amplification, slides are DAPI stained for nuclei then placed in the Xenium Analyzer with run reagents following CG000584. The gene panel is uploaded to the Xenium Analyzer and a primary image is take of the slides. All areas with visible tissue are selected for imaging and the imaging run is started. The Xenium processes imaging data during the run, identified cell boundaries and outputting image files along with cell by transcript by location matrices for further downstream analysis by Seurat 10x Xenium protocol. *Proximity Analysis of ARG1+ cells and IL6+ fibroblasts:* The output files of Xenium Analyzer were converted to an AnnData object that contained the cell centroid coordinates and raw transcript counts. A filter of 10 counts per cell and 5 cells per gene was applied to filter out low-quality cells and sparsely detected genes. Counts were normalized to 1e4 and log-transformed. Cells were then grouped into one of the following three categories: (1) ARG1+; (2) IL6+ARG1- and either CTHRC1 or COL1A1; and (3) IL6+ARG1- and CTHRC1 or COL1A1, with positive expression of a marker defined as having non-zero counts. *Co-occurrence probability ratio:* The ‘Analyze Xenium data’ tutorial in Squidpy (version 1.5.0) (61) was followed to compute the co-occurrence probability ratio of ARG1+ and fibroblasts as defined above. We performed 50% random subsampling of the data 5 times to compute the statistic and generate confidence intervals bounded by the minimum and maximum probability ratio generated by the subsamples. The co-occurrence probability ratio was computed at 25 μm between cells. Using the implementation in the 1.5.0 release of Squidpy, for each radial distance, the ratio of cells belonging to category exp (exp being any of the three categories of cell defined above) out of all cells within the given distance of an ARG1+ cell was averaged across all ARG1+ cells to compute the conditional probability in the numerator P(exp | ARG1+), while the denominator P(exp) was computed similarly but averaging probabilities that were computed by centering around every cell in the sample. Previous releases of Squidpy implemented a version of this test in the function gr.co_occurrence that used discrete interval bins (only including cells within two consecutive choices of radial distances). We chose to use inclusive intervals (including all cells within a given radial distance) for a more robust calculation of the co-occurrence probability ratio. We implemented our method into the codebase of Squidpy, with the help of Squidpy’s authors, and it is now the default implementation of the function gr.co_occurrence, starting in release 1.5.0.

### Analysis of published datasets

*Bleomycin time course:* Time series of single-cell data after bleomycin lung injury were obtained from Tsukui et al (8) and Strunz et al (9). These samples were processed and merged with Seurat (4.3.0) using the function SCTransform to correct for batch effects and annotated with SingleR 1.10.044 using the ImmGen database as reference to identify cell types. *Macrophage Annotation in vivo:* We annotated macrophages from Strunz et al. (9) according to the macrophage subtypes (C1, C2, C3) defined by Aran et al (2). and by Li et al (11) using SingleR(2). *BAL Microarray:* Microarray RNA data from bronchoalveolar cells of healthy individuals and IPF patients was extracted from the GPL14550 dataset within the GSE70867(12) repository. The differentially expressed genes between IPF and healthy samples were obtained by following the R workflow provided by the NCBI GEO2R platform using GEOquery (2.66.0) and limma (3.54.2) packages. *Gene expression of cultured mouse lung-derived macrophages:* The top markers of macrophage subtypes as defined by Li et al. (11) and Aran et al.(2) were quantified in our CD11b+, CSF1-treated macrophages (from the WT-WT coculture condition). *Analysis of lung tumor neutrophils:* A high-resolution single-cell atlas of tumor-associated neutrophils in non-small cell lung cancer was obtained from (32). The scANVI algorithm-based integrated NSCLC transcriptome atlas provided more than 1.2M cells from 19 studies and 309 patients. The metadata embedded with Leiden-clustering and cell-type annotations were utilized to identify and subset neutrophils. The subset data was log transformed and scaled, followed by filtering for cells expressing both ARG1 and IL6R. Pearson correlation coefficient, p-value and R-squared values were calculated for the correlation between ARG1 and IL6R.

### Ornithine Measurement

Cultured PCLS from IPF explanted lungs lysed in water, IPF lung samples lysed in RIPA buffer, or mouse lung homogenized and lysed in RIPA buffer were used to measure ornithine following manufacturer’s protocol (IS I-1000R, Immusmol). Briefly, each sample was precipitated with 50 µL precipitating reagents (provided by manufacturer) followed by extraction using D-reagent (a derivatizing agent provided by manufacturer) for 2h at room temperature. 10 µL of the prepared standards, controls and samples were incubated with 90 µL of antiserum overnight at 4DC. Subsequently, plates were washed and briefly incubated with enzyme conjugate for 30min followed by addition of substate. Absorbance was measured using microplate reader at 450nm.

### LC-MS metabolomics

Primary murine lung macrophages and fibroblasts were cocultured with or without 1 μM CB-1158. Fibroblasts were isolated by antibody-coated bead-based negative selection of macrophages. Fibroblast extracts were used to calculate protein equivalents by resuspension in 0.2 M NaOH, heated to 95 °C for 25 min, and determined via BCA (Pierce, 23225). Dried metabolites were resuspended in 50% ACN:water and 1/10th of the volume was loaded onto a Luna 3 um NH2 100A (150 × 2.0 mm) column (Phenomenex). The chromatographic separation was performed on a Vanquish Flex (Thermo Scientific) with mobile phases A (5 mM NH_4_AcO pH 9.9) and B (ACN) and a flow rate of 200 µL/min. A linear gradient from 15% A to 95% A over 18 min was followed by 9 min isocratic flow at 95% A and re-equilibration to 15% A. Metabolites were detected with a Thermo Scientific Q Exactive mass spectrometer run with polarity switching (+3.5 kV / −3.5 kV) in full scan mode with an m/z range of 65-975. TraceFinder 4.1 (Thermo Scientific) was used to quantify the targeted metabolites by area under the curve using expected retention time and accurate mass measurements (< 5 ppm). Values were normalized to cell number and sample protein concentration. Relative amounts of metabolites were calculated by summing up the values for all isotopologues of a given metabolite.

### Study Approval

Experiments in mice were performed in accordance with approved protocols by the University of California, San Francisco Institutional Animal Care and Use Committee. The studies with human tissues described in this paper were conducted according to the principles of the Declaration of Helsinki. Written, informed consent was obtained from all subjects, and the study was approved by the University of San Francisco, California and Vitalant institutional review board. With respect to deceased donor control explanted lungs, because samples were acquired from deceased individuals the study is not considered human subjects research as per UCSF and NIH policy.

### Statistics

One-way ANOVA followed by post hoc Sidak’s multiple comparisons tests was used for comparisons among more than two groups, and Student’s t-test was used for comparison between two groups. A p value less than 0.05 was considered significant. Analysis appropriately corrects for multiple comparisons and repeated measures.

### Data availability

Newly generated single-cell sequencing data will be made public on acceptance for publication at GSE242510.

## Declaration of Interests

The authors have declared that no conflict of interest exists.

## Acknowledgements

This work was supported by funding from the US Department of Defense (Grant W81XWH2110417 to MB), the UCSF Bakar Aging Research Institute (Investigator Grant to MB), Longevity Impetus Grants from Norn Group (MB and JGO), a UCSF Bakar Aging Research Institute investigator award (MB), the Nina Ireland Program for Lung Health Innovative Grant Program (PY), and NIH (1R01HL142701-01 to ABM and R01AI172754 SKP). The authors would like to acknowledge the staff within the Biological Imaging Development CoLab (BIDC) at UCSF Parnassus Heights, particularly Kyle Marchuk and Austin Edwards, for their support in microscopy experiments. Sequencing performed at the UCSF CAT was supported by UCSF PBBR, RRP IMIA, and NIH 1S10OD028511-01. We are grateful to Dr. Clifford Lowell for his valuable comments and suggestions on the manuscript. We thank Giovanni Palla and Nathan Levy for help in extending the Squidpy package.

## Author contributions

PY performed or assisted in all the experiments and in figure preparation. JGO performed single cell sequencing experiments and analysis and figure preparation. KC and SP performed single cell library preparation and sequencing, supervised by ARA. NB and AB assisted in functional experiments with lung tissues and cells. PD assisted in human neutrophil isolation and coculture experiments under the supervision of SKP. XY assisted PY in coculture experiments and microscopy under the supervision of BL. JN performed flow cytometry of mouse lung cells supervised by ABM. KB, TJ, and JW performed mouse breeding and development of crosses for experiments. TT and DS assisted in analysis of single cell sequencing data from mouse lung fibroblasts. MM and HT provided deceased donor control lung tissues supervised by MAM. LM, AG, CM, and VA performed Xenium spatial transcriptomic analysis supervised by WE. CC and AW designed and implemented spatial proximity analysis of Xenium data. RM generated and provided P2x4 floxed mice. WT and ST performed MIBI analysis under the guidance of TB. PJW provided lung explants from IPF patients and helped to interpret associated imaging data. KMT designed and performed metabolomic assays, assisted by PY. MB conceived of the work, supervised experimental planning and execution, and wrote the manuscript with input from PJW and KMT.

## Notes

### Competing Interest Statement

The authors have declared no competing interest.

### Summary of Updates

Further data have been added to address human/clinical/translational relevance. The paper is now expanded in number of figures to accommodate these data.

