## Supplementary Figures for "Myeloid-mesenchymal crosstalk drives Arg1-dependent profibrotic metabolism via ornithine in lung fibrosis"

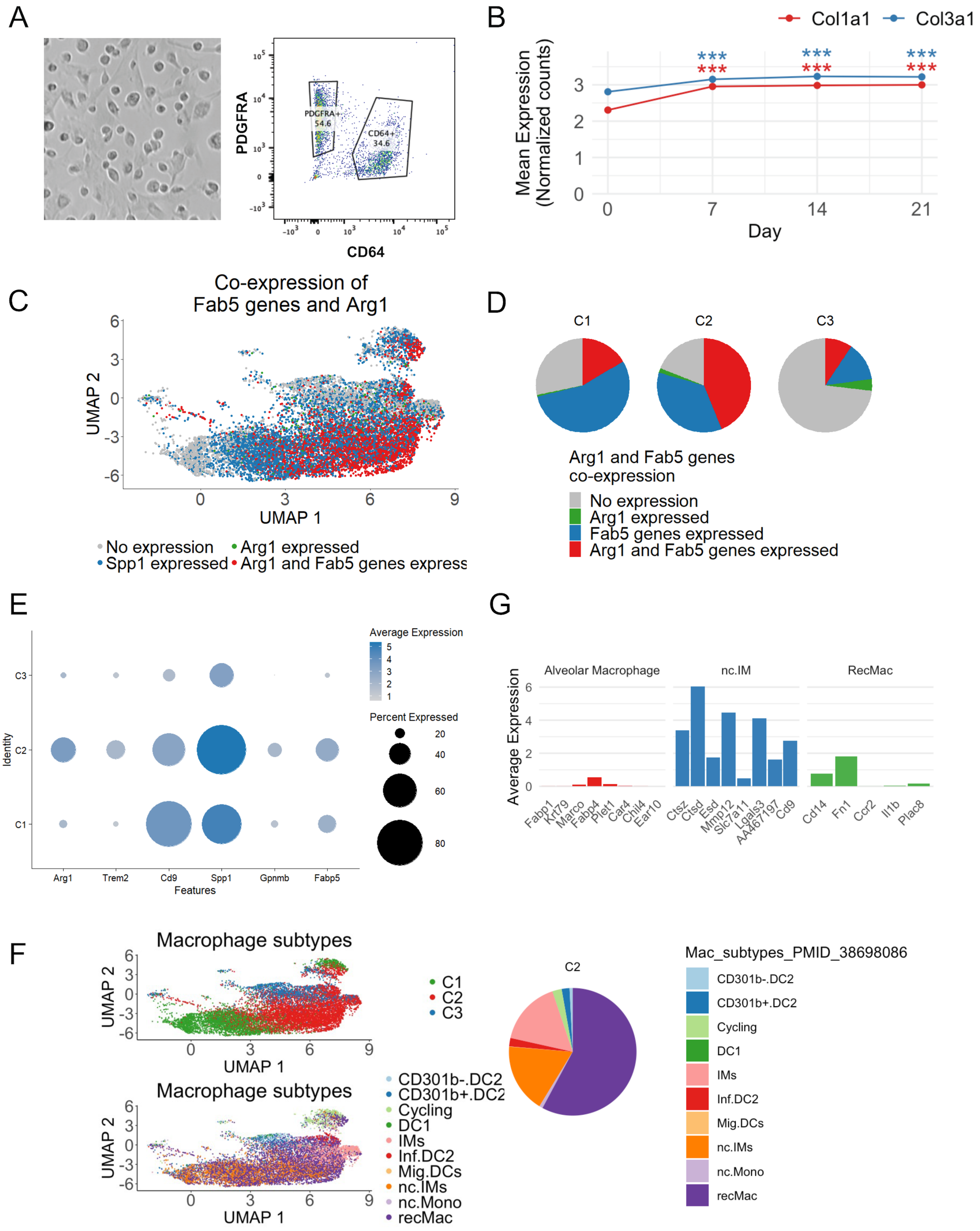


**Supplemental Figure 1**

A) Left: Bright field image of cocultured primary macrophages and fibroblasts. Right: Flow cytometry for Pdgfra and CD64 with coculture cells.
B) Expression of collagen genes across time after bleomycin injury in lung fibroblasts from Tsukui et al. (1) ***$p<0.001$.
C) Feature plot of lung macrophages from Strunz et al. (2) with Arg1 and Spp1 expression as well as Fab5 profibrotic macrophage genes from Fabre et al. (3) indicated.
D) Pie charts show proportions C1, C2, or C3 lung macrophages in (C). C1=alveolar macrophages; C2=transitional monocyte-derived macrophages; C3=monocyte derived macrophages; from Aran et al. (4).
E) Dot plot showing lung macrophage profibrotic genes from Fabre et al. (3) in macrophages from Strunz et al. (2) annotated as either C1, C2, or C3 macrophages
F) Annotation of cells from (C) by transcriptomic comparison using SingleR(4) to interstitial macrophage subtypes form Li et al. (2).
G) Analysis of expression of macrophage marker genes in the scRNAseq data for our CD11b+, CSF1-treated cultured macrophages (from the WT-WT coculture condition) to determine their likeness to macrophage subtypes defined by marker genes for interstitial macrophages from Li et al. (5) (RecMac and nc.IM), or for alveolar macrophages from Aran et al. (4).

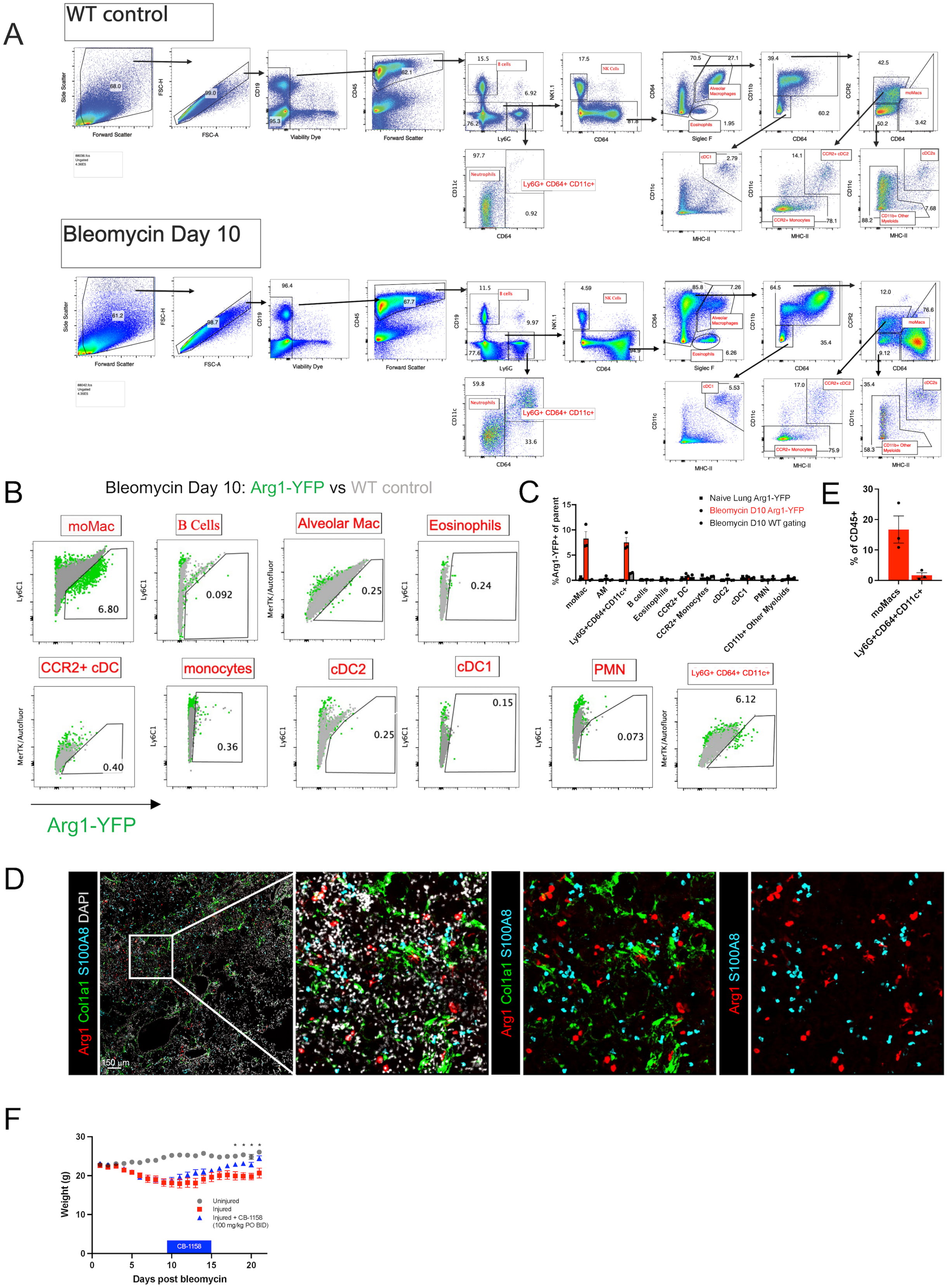


**Supplemental Figure 2**

A) Gating strategy for flow cytometry of lung cells at steady state. AM=alveolar macrophage (CD64+SiglecF+). MoMac=monocyte-derived macrophage (CD64+CD11b+SiglecF-).

B) Representative flow cytometry of mouse lungs from Arg1-YFP reporter and WT mice at steady state and 10 days post-bleomycin.

C) Plot of percentage of cells expressing YFP in each lineage, from flow cytometry corresponding to representative gating shown in (A) and (B). N=3 mice per condition. +/- SEM.

D) Lung immunofluorescence for neutrophil marker S100A8 from Arg1-RFP-CreERT2: R26-LSL-TdTomato : Col1a1-GFP mice, with no overlap between S100A8 and Arg1 detected. Data is representative of N=3 mice.

E) Plot of percentage of CD45+ cells accounted for by moMacs and Ly6G+CD64+CD11c+ cells at day 10 post-bleomycin, from flow cytometry corresponding to representative gating shown in (B). N=3 mice per condition. +/- SEM.

F) Weight measured daily in CB-1158-treated mice (corresponding to Figure 3a). *$p<0.05$ by Student's t-test. +/- SEM.

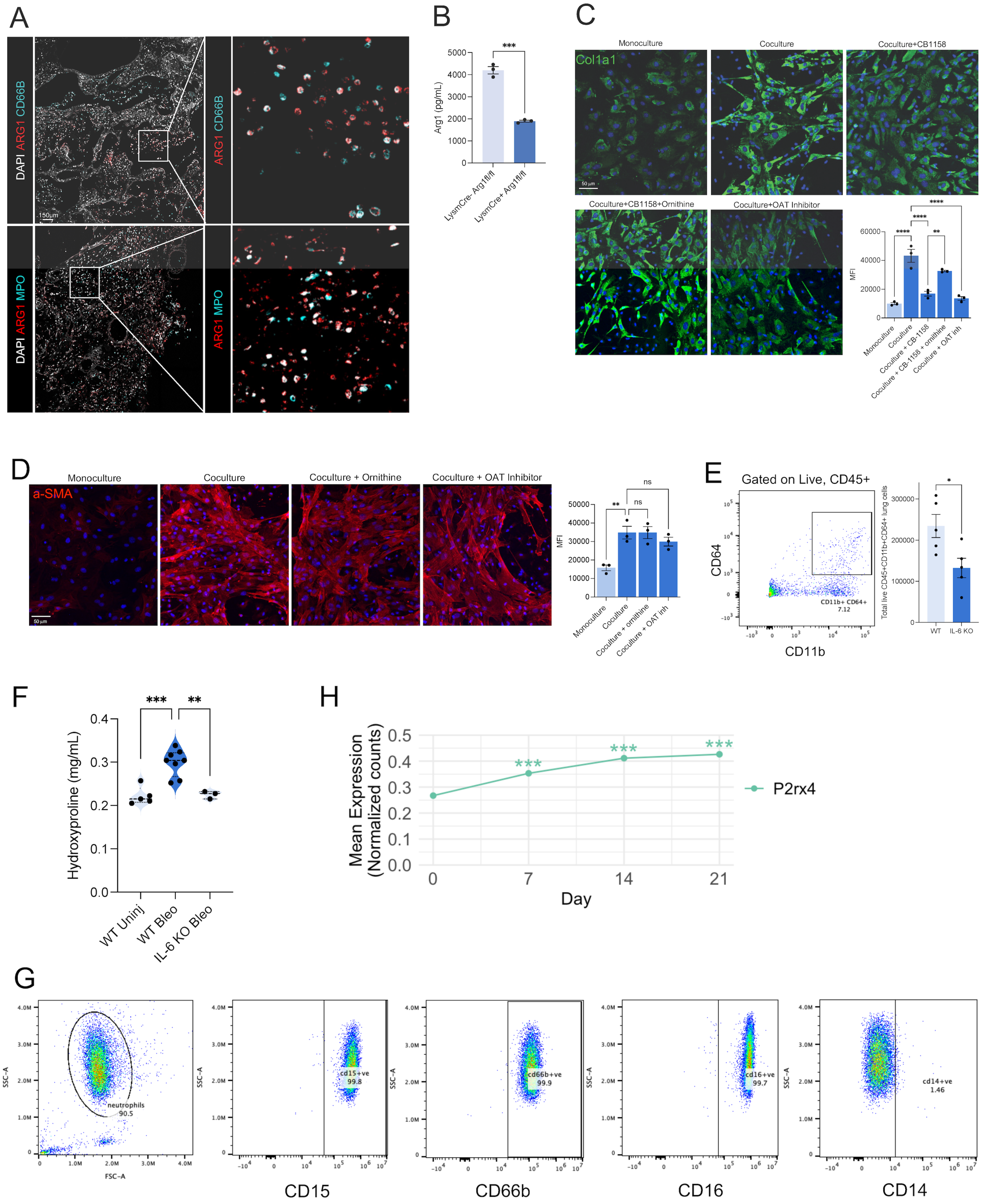


**Supplemental Figure 3**

A) Immunofluorescence of precision-cut lung slices (PCLS) from IPF lung. Displayed photomicrograph is representative of N=4 replicates.

B) Arg1 ELISA of lysates of CD11b+ cells freshly isolated by CD11b affinity column ***p<0.001 by Student's t test. Quantitation is for N=3 mice per condition. +/- SEM.
C) Col1a1 immunofluorescence of monocultured mouse lung fibroblasts with or without ornithine, Arg1 inhibitor CB-1158, or OAT inhibitor treatment. Quantification is for n=3 separate cultures each. **p<0.01, ****p<0.000 by 1-way ANOVA followed by post hoc Sidak's multiple comparisons tests. Bars show +/- SEM.
D) Alpha-smooth muscle actin immunofluorescence of monocultured mouse lung fibroblasts or murine lung macrophage-fibroblast cocultures, with or without ornithine or OAT inhibitor treatment. Quantification is for n=3 separate cultures each. **p<0.01 by 1-way ANOVA followed by post hoc Sidak's multiple comparisons tests. Bars show +/- SEM.
E) Representative FACS gating and quantitation of absolute lung cell number of live, CD45+CD11b+CD64+ cells in WT and IL-6 KO mice 14 days after bleomycin injury. Quantitation is for n=5 mice in each condition. *p<0.05 by Student's t test. Bars show +/- SEM.
F) Lung hydroxyproline for WT and IL-6 KO mice at 21 days after bleomycin injury. N=5, 8, 3 mice per condition, left to right. **p<0.01, ***p<0.001 by 1-way ANOVA followed by post hoc Sidak's multiple comparison's tests. Bars show +/- SEM.
G) Representative flow cytometry of neutrophils isolated from human donor peripheral blood. Individual markers are for the cells gated in the leftmost panel.
H) Expression of P2rx4 across time after bleomycin injury in lung fibroblasts from Tsukui et al.(1) ***p<0.001.
